# The fitness effects of codon composition of the horizontally transferred antibiotic resistance genes intensify at sub-lethal antibiotic levels

**DOI:** 10.1101/2023.02.03.526952

**Authors:** Michael Shaferman, Melis Gensel, Noga Alon, Khawla Alasad, Barak Rotblat, Adrian W.R. Serohijos, Lital Alfonta, Shimon Bershtein

**Affiliations:** Department of Life Sciences, Ben-Gurion University of the Negev, Beer-Sheva, Israel; Departement de Biochimie, Universite de Montreal, Quebec, Canada; National Institute for Biotechnology in the Negev (NIBN), Beer-Sheva, Israel; Centre Robert-Cedergren en Bio-informatique et Genomique, Universite de Montreal, Quebec, Canada; Department of Chemistry, Ben-Gurion University of the Negev, Beer-Sheva, Israel

## Abstract

The rampant variability in codon bias existing between bacterial genomes is expected to interfere with horizontal gene transfer (HGT), a phenomenon that drives bacterial adaptation. However, delineating the constraints imposed by codon bias on functional integration of the transferred genes is complicated by multiple genomic and functional barriers controlling HGT, and by the dependence of the evolutionary outcomes of HGT on the host’s environment. Here, we designed an experimental system in which codon composition of the transferred genes is the only variable triggering fitness change of the host. We replaced *E. coli*’s chromosomal *folA* gene encoding dihydrofolate reductase, an essential enzyme that constitutes a target for trimethoprim, with combinatorial libraries of synonymous codons of *folA* genes from trimethoprim-sensitive *Listeria grayi* and trimethoprim-resistant *Neisseria sicca.* The resulting populations underwent selection at a range of trimethoprim concentrations, and the ensuing changes in variant frequencies were used to infer the fitness effects of the individual combinations of codons. We found that when HGT causes overstabilization of the 5’-end mRNA, the fitness contribution of mRNA folding stability dominates over that of codon optimality. The 5’-end overstabilization can also lead to mRNA accumulation outside of the polysome, thus preventing the decay of the foreign transcripts despite the codon composition-driven reduction in translation efficiency. Importantly, the fitness effects of mRNA stability or codon optimality become apparent only at sub-lethal levels of trimethoprim individually tailored for each library, emphasizing the central role of the host’s environment in shaping the codon bias compatibility of horizontally transferred genes.

## Introduction

The genetic code is redundant. With the exception of methionine and tryptophan, each amino acid is encoded by two to six codons. The synonymous codons are not used with equal frequencies, resulting in codon usage bias (Grantham, Gautier and Gouy 1980; Grantham, Gautier, Gouy, et al. 1980; Gouy and Gautier 1982; Sharp and Li 1987; Chen, et al. 2004; Tuller, Carmi, et al. 2010; Goodman, et al. 2013; Pechmann and Frydman 2013; Chaney, et al. 2017; Jacobs and Shakhnovich 2017). The codon bias observed among different genomes is thought to be driven predominantly by genome-wide mutational (*i.e.*, neutral) processes that shape the global nucleotide composition, rather than by selection on individual coding sequences (Knight, et al. 2001; Chen, et al. 2004). In contrast, codon bias found *within* genomes cannot arise from neutral processes alone, and requires the involvement of selection (Hershberg and Petrov 2008). The selectionist view of the intragenomic codon bias is corroborated by three robust phenomena detected across multiple prokaryotic and eukaryotic species. The first phenomenon is the universal reduction in the folding stability of the 5’-end mRNA secondary structure found in nearly all species tested, which is interpreted to be a result of selection pressure to optimize translation initiation (Kudla, et al. 2009; Gu, et al. 2010; Tuller, Waldman, et al. 2010; Bentele, et al. 2013; Goodman, et al. 2013; Bhattacharyya, et al. 2018). The second phenomenon is the match between frequently used codons and abundant tRNA molecules (Ikemura 1985; Yamao, et al. 1991; Moriyama and Powell 1997; Kanaya, et al. 1999; Duret 2000; Kanaya, et al. 2001; Rocha 2004). Selection may favor more frequent codons because optimizing the speed and accuracy of translation elongation increases the availability of free ribosomes and reduces the frequency of translational mutations (Gouy and Gautier 1982; Duret and Mouchiroud 1999; Ghaemmaghami, et al. 2003; Goetz and Fuglsang 2005; Plotkin and Kudla 2011). The third phenomenon is the association of non-optimal (less frequent) codons with the formation of co-translational folding intermediates (Pechmann and Frydman 2013; Jacobs and Shakhnovich 2017). Reduction in the rates of translation elongation induced by clusters of non-optimal codons is thought to be beneficial for promoting the formation of native protein structures on the ribosome (Komar 2009; Schlesinger, et al. 2017).

An important outcome of the neutral and adaptive forces operating on synonymous codons is the emergence of distinct patterns of codon bias in the genomes of different organisms (Deng, et al. 2020). Bacterial genomes, in particular, exhibit stark differences in both genome-wide usage of codons and in non-random distribution of codons within individual genes (Bentele, et al. 2013). Bacterial genomes are also continuously reshaped via gene loss and gene acquisition (Popa and Dagan 2011; Soucy, et al. 2015). Genes are acquired by bacteria in a process dubbed ‘horizontal gene transfer’ (HGT). HGT has the potential to induce rapid and profound phenotypic changes and is considered to be a major driving force of bacterial adaptation and speciation (Ochman, et al. 2000; Nakamura, et al. 2004). HGT, however, may be hindered by codon bias variability among bacterial genomes. Indeed, when a foreign gene integrates into the genome of a bacterial host, the codon incompatibility between the donor and the recipient may cause an inadequate expression of the newly acquired gene and/or incur a fitness cost on the host (Amoros-Moya, et al. 2010; Lind, et al. 2010; Bershtein, Serohijos, et al. 2015). It is, therefore, imperative to define the impact of codon bias on the establishment and maintenance of successful HGT events in bacteria.

The task of delineating the role of codon bias in HGT is complicated by multiple genomic and functional barriers that constrain HGT (Thomas and Nielsen 2005; Lind, et al. 2010; Popa and Dagan 2011; Bershtein, Serohijos, et al. 2015; Kacar, et al. 2017), and by dependence of the evolutionary outcomes of HGT on a particular environment of the host (Amoros-Moya, et al. 2010). Computational studies that analyzed established HGT events among bacterial genomes revealed that HGT frequency positively and strongly correlates with the similarity of tRNA pools between donors and acceptors, suggesting that the compatibility between codon usage of the donor and tRNA pools of the host, *i.e.*, codon optimality, constitutes an important codon usage-related requirement to successful HGTs (Medrano-Soto, et al. 2004; Tuller, et al. 2011). Such analyses, however, do not reveal the molecular mechanisms underlying failed instances of HGT. Indeed, Kudla *et*. *al*. have demonstrated that only 5% of the variability in fluorescence of synthetic GFP genes with diversified synonymous codons expressed in *E. coli* can be explained by codon optimality measured by Codon Adaptation Index (CAI), whereas the folding stability of mRNA secondary structure near the translation start codon explained almost 60% of the fluorescence variability (Kudla, et al. 2009). Conversely, higher CAI values of GFP variants correlated significantly with faster bacterial growth, possibly by sequestering less ribosomes than variants carrying low-frequency codons, whereas no correlation was observed between the 5’-end mRNA folding stability and growth. These findings provided a clear first indication that both mRNA folding stability at the starts of genes and codon optimality constitute barriers to the heterologous gene expression. However, mRNA stability and codon optimality cannot be correlated to the fitness of cells carrying individual GFP variants since GFP function is irrelevant to organismal physiology. A direct link between the function of the transferred genes and fitness of the host was established in another experimental study that measured the effects of two hundred diverse antibiotic resistance-conferring plasmid genes on *E. coli* growth (Porse, et al. 2018). Surprisingly, no significant contribution of the mRNA folding stability or codon usage preferences to the compatibility of the horizontally transferred genes was detected. Instead, resistance mechanisms were deemed to be major factors that control the compatibility of the heterologous antibiotic resistance genes (Porse, et al. 2018). It is reasonable to conclude that the very high antibiotic concentrations used in this study (up to 30-fold the minimal inhibitory concentration (MIC)) potentially minimized the effect of sequence composition on fitness. This notion is particularly relevant, since bacteria in nature and in clinic are often exposed to non-lethal (sub-MIC) levels of antibiotics (Gullberg, et al. 2011; Hughes and Andersson 2012). Further, the inferring of the contribution of codon bias of the various antibiotic resistance genes to *E. coli*’s growth could have been masked by mechanisms affecting the abundance and function of the protein products of the transferred genes downstream to transcription and translation steps, and, therefore, being unrelated to codon bias. These mechanisms may include contrasting sensitivities of transferred proteins to degradation and aggregation within the milieu of the host cell(Bershtein, Serohijos, et al. 2015), uneven efficiencies in protein transport to the periplasm or differences in fitness cost associated with such a transfer (Rajer and Sandegren 2022), or the formation of spurious interactions with cellular counterparts of the host (Bhattacharyya, et al. 2016).

We reasoned that the incongruities found between the studies discussed above can be settled by establishing a clear link between codon bias of the transferred genes and fitness of the host. To this end, we used *folA* gene encoding dihydrofolate reductase (DHFR), an essential enzyme that constitutes a target for trimethoprim (TMP), as a model for HGT. We replaced the open reading frame of *E. coli*’s chromosomal copy of the *folA* gene (while preserving the endogenous regulatory region) with combinatorial libraries of synonymous codons of *folA* genes from TMP-sensitive *Listeria grayi* and TMP-resistant *Neisseria sicca*. We found that both mRNA stability at the translation initiation regions and codon optimality impact the fitness of the host, however this effect is prominent only within a relatively narrow range of sub-MIC TMP concentrations individually tailored for each xenologue, underlining the importance of the host’s environment in shaping the codon bias compatibility of horizontally transferred genes.

## Results

### *folA* gene as a model of HGT

To tease apart the constraints imposed by codon bias on the functional integration of horizontally transferred genes, we devised the following experimental criteria. First, physiologically relevant changes in the function and abundance of the protein products of transferred genes should produce fitness effects in the host. Second, factors affecting the abundance and activity of foreign proteins downstream to the transcription and translation steps must not mask the changes in protein abundances caused by codon bias. Third, the fitness effects of the transferred genes must be measured at a range of suitable environmental conditions. To comply with these criteria, we performed an artificial horizontal gene transfer by replacing the open reading frame of the chromosomal *E. coli folA* gene encoding dihydrofolate reductase (DHFR) with various bacterial xenologoues. The first criterion is met in such an experimental setup through the essentiality of DHFR in *E. coli*. DHFR catalyzes NADPH-dependent reduction of dihydrofolate to tetrahydrofolate. The latter is an essential carrier for the one-carbon transfer reactions required for nucleic acids and amino acids metabolism (Schnell, et al. 2004). Changes in functional DHFR levels, including reduction in DHFR abundance induced by the diversification of synonymous codons (Bhattacharyya, et al. 2018), are known to directly control bacterial growth (Bershtein, et al. 2013; Bershtein, Choi, et al. 2015; Bershtein, Serohijos, et al. 2015; Bhattacharyya, et al. 2016; Rodrigues, et al. 2016). When the product of DHFR abundance and catalytic activity (*k_cat_*/K_M_) drops below the basal level, which in *E. coli* constitutes 40-100 DHFR molecules per cell (Taniguchi, et al. 2010), the decrease in bacterial growth rate follows Michaelis-Menten-like kinetics (Bershtein, et al. 2013; Bershtein, Choi, et al. 2015; Rodrigues, et al. 2016; Bhattacharyya, et al. 2018). The drop in functional DHFR levels was shown to cause a profound imbalance in the metabolic pools (Bershtein, Choi, et al. 2015; Bhattacharyya, et al. 2021), which, in turn, leads to an upregulation of *folA* transcription via a negative metabolic regulation loop operating through binding of TyrR transcription activator to two TyrR boxes located in *folA* regulatory region (Yang, et al. 2007). Furthermore, the increase in DHFR abundance, well above the basal level, was also found to cause a decline in the growth rate of *E. coli*, owing to the formation of transient protein-protein interactions that trigger toxic metabolic imbalance (Bhattacharyya, et al. 2016).

Upon the horizontal transfer of xenologous *folA* genes, changes in DHFR levels can occur not only as a result of codon bias, but also due to the differential sensitivities of the xenologous DHFR proteins to aggregation and degradation in the cytoplasm of *E. coli* (Bershtein, et al. 2013; Bershtein, Serohijos, et al. 2015). Thus, to comply with the second criterion, we measured and compared changes in fitness effects caused by codon usage diversification within each individual xenologous *folA* strain, rather than comparing fitness effects across strains.

To fulfill the third criterion, we took advantage of the DHFR sensitivity to TMP, a widely-used antibiotic that competitively binds to active sites of bacterial DHFRs (Burchall and Hitchings 1965). We measured the fitness effects incurred by codon bias of xenologous DHFR proteins on *E. coli* growth at a range of TMP concentrations, including sub-MIC levels.

Lastly, comparative genomics studies have demonstrated that HGT plays an important role in the evolution of folate metabolic pathway in bacteria, including the spread of antifolate resistance (de Crecy-Lagard, et al. 2007; Stern, et al. 2010), making DHFR a suitable target to study the constraints imposed by codon bias on HGT.

We assembled a collection of diverse xenologous *folA* genes from *Bacillus subtilis*, *Lactococcus lactis*, *Leuconostoc mesenteroides*, *Neisseria sicca*, *Methyloccocus capsulatus*, *Listeria grayi*, and *Pseudomonas putida* that differ with respect to their sequence composition (GC content ranges from 34% to 66%), evolutionary distance (between 31% to 49% amino acid sequence identity relatively to *E. coli*’s DHFR), and catalytic efficiency of their protein products (*k_cat_/K_M_* values of DHFRs range from 3.9 to 71.7 μM^-1^sec^-1^) (**table 1**, **table S1**).

**Table 1.** Xenologous folA genes

| # | Origin of *folA* gene |  |  |  |  |  |
| --- | --- | --- | --- | --- | --- | --- |
| | Species | Class | GC<sup>a</sup><sub>genome</sub>, % | GC<sup>b</sup><sub>*folA*</sub>, % | AA<sup>c</sup><sub>identity</sub>, % | $k_{cat}/K_M$<sup>d</sup>, $\mu\text{M}^{-1}\text{sec}^{-1}$ |
| 1 | *Bacillus subtilis* (B.su) | Bacilli | 44 | 43 | 44 | 30.8 |
| 2 | *Lactococcus lactis* (L.la) | Bacilli | 35 | 34 | 31 | 44.2 |
| 3 | *Leuconostoc mesenteroides* (L.me) | Bacilli | 38 | 42 | 32 | 28.1 |
| 4 | *Neisseria sicca* (N.si) | Betaproteobacteria | 51 | 58 | 44 | 62.5 |
| 5 | *Methylococcus capsulatus* (M.ca) | Gamma proteobacteria | 64 | 66 | 49 | 3.9 |
| 6 | *Listeria grayi* (L.gr) | Bacilli | 42 | 45 | 37 | 19.5 |
| 7 | *Pseudomonas putida* (P.pu) | Bacilli | 60 | 63 | 42 | 71.7 |
| 8 | *Escherichia coli* (E.co) | Gamma proteobacteria | 51 | 53 | 100 | 8.1 |

<sup>a</sup>GC content of the original genome (Bentele, et al. 2013)
<sup>b</sup>GC content of *folA* genes (see Table S1)
<sup>c</sup>% amino acid identity relatively to *E. coli*'s DHFR
<sup>d</sup>from ref (Bershtein, et al. 2013)

Using CAI and tAI codon usage metrics of the donor bacteria and *E. coli* (Sharp and Li 1987; Tuller, et al. 2011) (**table S2**), we calculated how the CAI and tAI values of the xenologous *folA* genes, CAI*g* and tAI*g* (defined as the geometric mean of the CAI or tAI of all codons of a gene), would change upon horizontal transfer to *E. coli*. CAI and tAI scales estimate how codons affect translational efficiencies. CAI assumes that frequent codons are used more often in highly expressed genes (Sharp and Li 1987). tAI relies on relative tRNA abundance and on constraints imposed by codon-tRNA wobble interactions (dos Reis, et al. 2004). We determined that the transfer of the xenologous *folA* genes to the chromosome of *E. coli* would be accompanied by a drop in CAI*g* and tAI*g* values for all the xenologues, indicating that *E. coli* features a rather different codon usage pattern than the chosen donor species (**fig. S1A,B**).

As mentioned above (Gu, et al. 2010), genes are universally subject to selection pressure to reduce mRNA secondary structure near the translation start codon. However, if a horizontally transferred gene is acquired without the original promoter, it must be inserted near a promoter of the host to be expressed. The combination of the 5’-untranslated regulatory region of the host with the synonymous codons near the start codon of the donor gene may induce changes in the folding stability of the secondary mRNA structure of a xenologue, and, potentially, affect the efficiency of translation. We, therefore, calculated the expected changes in the folding stability profiles (Gibbs free energy difference between unfolded and folded states of mRNA, Δ*G*) of the 5’-end mRNA transcripts of the xenologous *folA* genes in a scenario in which the original sequence immediately upstream to the protein coding region, as found in the donor organism, is replaced with the *folA* regulatory sequence of the host upon horizontal transfer to *E. coli*. Since *folA* transcription in *E. coli* commences at nucleotide −25, the mRNA folding stability profiles were calculated for 30-nt long fragments, the length that roughly corresponds to a ribosomal footprint size (Martens, et al. 2015), from nucleotide −25 of the upstream sequence and up to nucleotide +5 within the coding sequence (see **Materials and Methods**). As anticipated, we found a wide range of stability effects near the translation start site (**fig. S1C**). In some xenologues, mRNA transcripts exhibited a 1 to 3 kcal/mol increase in folding stability relative to the original organism, whereas in others loss of folding stability ranged from 1 to 11 kcal/mol. It is worth noting, however, that in some donor organisms *folA* genes are transcribed as part of a larger polycistronic transcript, often immediately downstream to the coding sequences of other genes (**fig. S2**). Therefore, the possible contribution of the 5’-end *folA* mRNA folding stability to translation efficiency in these bacteria may differ markedly from other organisms where, like in *E. coli*, *folA* is transcribed as a single gene from a unique promoter.

Previously, the replacement of rare synonymous codons of the endogenous *folA* gene in *E. coli* with the most frequent codons produced a reduction in bacterial growth (Bhattacharyya, et al. 2018). We, therefore, assumed that a similar replacement within xenologous *folA* genes will also cause changes in the growth of the host. To test this assumption, we synthesized two sets of protein coding sequences for each of the seven chosen xenologous *folA* genes. The first set is composed of the original sequences as they appear in the donor organisms (from here on ‘ORG’ genes) (**table S1**), and the second set consists of modified sequences, in which the original synonymous codons were replaced with the most frequent codons in the *folA* gene of *E. coli* (‘MOD’ genes) (**fig. S3** and **table S3**). Using the same tactic, we also generated a MOD sequence for the endogenous *folA* gene from *E. coli*. As expected, codon replacement led to an increase in CAIg and tAIg values (calculated using *E. coli*’s codon usage metrics, **table S2**) in all of the modified sequences as well as to an increase in the GC content in sequences whose original GC content was <60% (**fig. S4** and **table 1**). Codon replacement also induced profound changes in the mRNA folding stability profiles of MOD sequences. mRNA stability was calculated using a 30 nucleotide-long sliding window starting from the mRNA transcription start (nucleotide −25) in steps of 1 nt for a total of 100 windows (see **Materials and Methods**). As a rule, MOD sequences were found to be more stable than their ORG counterparts (**fig. S5**). However, the extent of the change varied between strains and between different locations within the same mRNA transcript. Lastly, we replaced the open reading frame of the chromosomal copy of the *folA* gene of *E. coli* with each of the synthesized genes, while preserving the endogenous upstream regulatory region of *folA*, and measured the ensuing effects on the endogenous *folA* promoter activity, mRNA abundance, intracellular DHFR levels, and growth rates of the host either in the absence or presence of TMP.

### The original to frequent codon replacements within xenologous *folA* genes affects the strains’ sensitivity to sub-MIC trimethoprim levels

In the absence of TMP, codon replacement resulted in a 5%-20% decrease in the growth rates of four out of seven xenologous strains (*L. mesenteroides*, *N. sicca*, *M. capsulatus*, and *P. putida*), a trend similar to that observed upon rare to frequent codon replacement in the endogenous *folA* coding sequence in *E. coli* (**fig. 1A** and ref.(Bhattacharyya, et al. 2018)). In these strains, the drop in growth was matched by a decrease in the intracellular soluble DHFR fraction (**fig. 1B**). In the strain carrying *L. grayi*, however, a drop in protein abundance upon codon replacement was accompanied by a 30% *increase* in growth, suggesting that higher DHFR levels produced by the original sequence are toxic to *E. coli* (**fig. 1A,B**). In *L. lactis* strain, a drop in DHFR levels produced almost no effect on growth. No change in either growth rate or protein abundance was observed in the strain carrying modified *B. subtilis folA* gene (**fig. 1A,B**). The variability in the growth effects did not correlate with changes in CAIg/tAIg values or the GC content of the modified sequences. However, changes in tAIg values did correlate weakly, yet significantly, with changes in *folA* promoter activity, which was measured using a reporter plasmid in which the endogenous *folA* promoter is fused to GFP (Zaslaver, et al. 2006) (Pearson’s correlation R = 0.84, *p-value* = 0.02) (**fig. 1D** and **fig. S6A**). The correlation is inverse, meaning that higher tAIg values correlate with less active promoter. Following a similar trend, albeit below the threshold of significance, there was an observed correlation with CAIg - promoter activity (**fig. S6B**). CAIg/tAIg values are measures of codon optimality, and an increase in these values is expected to be associated with more efficient translation.

**Fig. 1.**
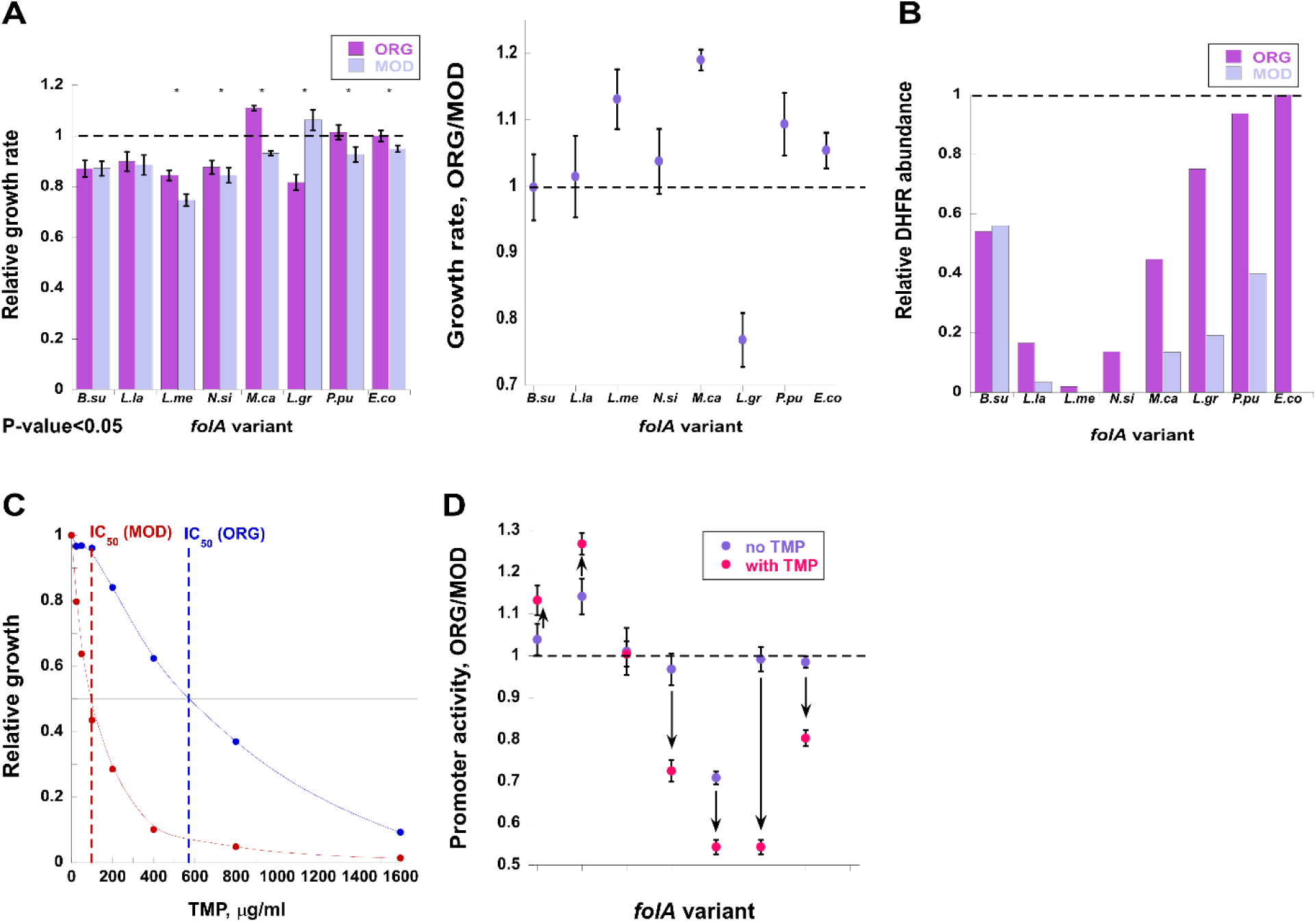
*C*odon replacements within *folA* xenologoues induce phenotypic effects in *E. coli* host. *folA* variants are marked as in **Table 1**. (*A*) *Left panel*. Growth rates of strains carrying original (ORG) and modified (MOD) *folA* sequences relative to growth of WT *E. coli* strain (dashed line). Asterisks indicate variants exhibiting significant differences in growth rates (two-tailed test, *p-value* < 0.05). *Right panel*. Growth ratios between ORG and MOD *folA* variants. Variants above the dashed line exhibit better growth with original *folA* sequence. (*B*) Intracellular DHFR abundance in strains carrying ORG and MOD *folA* sequences relative to DHFR abundance in WT *E. coli* (dashed line). (*C*) Normalized growth of *E. coli* strain carrying ORG (blue) or MOD (red) *folA* gene from *N. sicca* is plotted as a function of TMP concentrations. Dashed line intersections with the black line indicate TMP concentrations at which growth of each variant reaches 50% of its growth in the absence of TMP (IC50). (*D*) Changes in *folA* promoter activity upon codon replacement. Ratios of promoter activity between strains carrying ORG and MOD *folA* sequences are plotted for each variant in the absence of TMP (blue) and in the presence of IC50 amounts of TMP (red) (See also **table 2**). The direction of change upon TMP addition is indicated with black arrows. In variants found below the dashed line, the *folA* promoter tends to be more active with MOD *folA* sequences. Error bars were calculated from four independent measurements.

The correlation between a decrease in *folA* promoter activity, which is indicative of an increase in the functional DHFR levels (Bollenbach, et al. 2009; Bershtein, Choi, et al. 2015), with an increase in tAIg/CAIg values indeed indicates an increase in translation efficiency. Using a 30 nucleotide-long sliding window starting from the mRNA transcription start (25 nt upstream of the translation start codon) and moving 1 nt at a time, we also calculated the difference in the mRNA folding stability between the ORG and MOD *folA* xenologs, *ΔΔ*G. Changes in the promoter activity upon codon replacement correlated weakly, yet significantly, with mRNA *ΔΔ*G values near the translation start codons (windows 1 to 5) (**fig. S6E**). An increase in mRNA stability was accompanied by a stronger promoter activation, indicating that overstabilization of the 5’-end of mRNA structure interfered with translation and led to a reduction in functional DHFR levels (Spearman’s correlation *r* = 0.79, *p-value* = 0.035) (**fig. S6E**). However, no significant correlation between mRNA *ΔΔ*G values and changes in growth was observed.

To estimate the fitness effects of codon diversification in the presence of TMP, we exposed the xenologous strains to a wide range of TMP concentrations (below and up to MIC levels) and determined the concentrations that caused 50% drop in growth (IC50). With the exception of the *L. lactis* xenologous strain, which is highly resistant to TMP, growth differences between ORG and MOD strains became pronounced in the presence of sub-MIC levels of TMP, with ORG over MOD ratios of IC50 concentrations ranging between 0.44 to 5.8-fold for the xenologous strains and 18-fold for the endogenous *folA* from *E. coli* (**fig. 1C**, **fig. S7**, and **table 2**). In contrast, at TMP levels approaching a MIC, the difference in growth between strains carrying ORG and MOD *folA* sequences largely dwindled (**fig. 1C**, **fig. S7**, and **table 2**).

**Table 2.** MIC and IC50 concentrations of TMP for xenologous and endogenous E. coli strains carrying original (ORG) and modified (MOD) folA sequences.

| # | Origin of *folA* gene | ORG<sup>a</sup> |  | MOD<sup>b</sup> |  | IC<sub>50</sub><sup>e</sup> |
| --- | --- | --- | --- | --- | --- | --- |
|  |  | MIC<sup>c</sup> | IC<sub>50</sub><sup>d</sup> | MIC | IC<sub>50</sub> | ORG/MOD |
| 1 | *Bacillus subtilis* | >10 | 3.1 | >10 | 7.1 | 0.44 |
| 2 | *Lactococcus lactis* | >2,000 | 1,750 | >2,000 | 1,750 | 1 |
| 3 | *Leuconostoc mesenteroides* | >1,600 | 380 | >1,600 | 120 | 3.2 |
| 4 | *Neisseria sicca* | >1,600 | 580 | >1,600 | 100 | 5.8 |
| 5 | *Methylococcus capsulatus* | >10 | 5.8 | >10 | 3 | 1.9 |
| 6 | *Listeria grayi* | >2.5 | 0.37 | 3 | 0.1 | 3.7 |
| 7 | *Pseudomonas putida* | >1,600 | 1,200 | 1,600 | 590 | 2 |
| 8 | *Escherichia coli* | >10 | 1.8 | 2 | 0.1 | 18 |

<sup>a</sup>Strains carrying original *folA* sequences.
<sup>b</sup>Strains carrying original *folA* sequences.
<sup>c</sup>Minimal Inhibitory concentration of TMP, $\mu\text{g/ml}$ .
<sup>d</sup>TMP concentration inhibiting 50% of growth, $\mu\text{g/ml}$ .
<sup>e</sup>Ratio of IC<sub>50</sub> TMP concentrations between strains carrying ORG and MOD *folA* genes.

As a general trend, the replacement of original to frequent codons resulted in an increased sensitivity to sub-MIC levels of TMP. This was true also for the *L. lactis* strain, which, in the absence of TMP, showed an increase in growth of the strain carrying MOD sequence (**fig. 1A**). It is plausible that in the presence of TMP, a competitive inhibitor of DHFR, higher *L. lactis* DHFR abundance confers a fitness advantage. Conversely, in case of *B. subtilis* xenologous strain, the MOD sequence exhibited a reduced sensitivity to TMP (**table 2** and **fig. S7**). Interestingly, in contrast to other xenlogues, the 5’-mRNA region of the MOD sequences of *B. subtilis folA* exhibited almost 5 kcal/mol overstabilization compared to the ORG sequence (**fig. S5A**). Changes in CAIg/tAIg values or in GC content were not predictive of the growth effects observed in the host grown in the presence of TMP, but the observed changes in TMP sensitivity were largely captured by the response of the *folA* endogenous promoter (**fig. 1D**). Addition of TMP at IC50 levels led to an increase in the promoter activity in *N. sicca*, *M. capsulatus*, *L. grayi* and *P. putida* xenologous strains carrying MOD *folA* sequences. Conversely, in case of *B. subtilis* and *L. lactis* the promoter activity was markedly increased in strains carrying the ORG sequences, indicating that in these genes, codon replacement led to a reduction in TMP sensitivity. Overall, a weakly significant inverse correlation was observed between changes in tAIg values and changes in promoter activity (Pearson’s correlation R = 0.87, p-value = 0.01) (**fig. S6C**). Following a similar, albeit insignificant, trend an inverse correlation was between changes in CAIg values and promoter activity (**fig. S6D**). The increase in promoter activity upon codon replacement in the presence of sub-MIC TMP was also significantly correlated with mRNA thermodynamic stabilization near the translation start codon, similarly to what was observed in the absence of TMP (Pearson’s correlation R = 0.89, *p-value* = 0.007) (**fig. S6E**).

Collectively, these findings indicate that codon bias-related fitness effects vary substantially between the xenologues. Importantly, these effects are highly dependent on TMP concentrations. In fact, for each xenologue, a relatively narrow range of sub-MIC TMP concentrations exists in which the growth effects of codon replacement become pronounced, revealing the high sensitivity of codon bias-related fitness effects of the horizontally transferred antibiotic resistance genes to the environmental levels of antibiotics. The changes in CAIg/tAIg values, GC content, or 5’-end mRNA folding stability upon codon replacement were not predictive of the changes in growth of the xenologous strains. However, changes in both codon optimality (CAIg/tAIg values) and 5’-end mRNA thermodynamic stability were predictive of the changes in *folA* promoter activity.

### Competition among xenologous *folA* variants with diversified codons intensifies at sub-lethal TMP levels

Although it is clear from the obtained data that ORG to MOD codon replacements within xenologous *folA* strains directly affect *folA* promoter activity, intracellular DHFR abundance, host growth, and TMP sensitivity, it was not possible to reliably infer the changes in bacterial growth from changes in codon composition (CAIg/tAIg values or GC content) or mRNA stability, most plausibly because of the small number of xenologous sequences and a rather limited variety of codon replacements (only a single combination of replaced codons was tested). To address this problem, we chose to focus on *folA* genes from *N. sicca* and *L. grayi.* These xenologues differ substantially in their sensitivity to TMP: MIC of TMP in the *E. coli* strain carrying ORG *N. sicca folA* is around 1600 μg/ml TMP but only 2.5 μg/ml TMP in case of ORG *L. grayi folA* (**table 2**, **fig. 1C**, and **fig. S7D**). To identify the areas in the coding sequences of both genes that are responsible for the most pronounced codon bias-related fitness effects, we performed replacements of the fully modified sequences (MOD) back to ORG codons in stretches of 30 codons targeting the beginning, middle and end of both genes. The resulting chimeric, original, and fully modified strains were grown in the presence of sub-MIC TMP concentrations, and their fitness was compared. We determined that ≥ 80% of codon bias-driven fitness effects are localized to codons 1-30 (**fig. S8A,B**). Similar results were obtained when the MOD endogenous *E. coli folA* sequence was restored back to the ORG state (**fig. S8C**). Having determined that the observed codon bias-driven fitness effects are localized primarily within the first 15-30 codons, we generated 6 chimeric libraries, two for each of the *folA* sequences from *N. sicca*, *L. grayi*, and *E. coli*. To this end, the synonymous codons found within codons 1-15, or codons 16-30 of the fully modified *folA* sequences (MOD) were diversified back to the original codons in a combinatorial manner (**fig. 2B**). The resulting chimeric gene libraries were used to generate libraries of chimeric xenologous strains by replacing the chromosomal *folA* coding sequence of *E. coli* without perturbing the endogenous regulatory region upstream to the coding sequence (**fig. 2A**). Deep sequencing revealed that the generated chimeric strains covered between 55% to 98% of the theoretically possible combinations of codons (**fig. 2C**). To limit the possibility for occurrence of *de novo* mutations, the obtained libraries were subjected to only two serial passages, each 16 hours-long, at a range of sub-MIC TMP concentrations. At each time point (day0, day1, and day2), the libraries were analyzed by deep sequencing of the *folA* gene (**fig. 2A,C**).

**Fig. 2.**
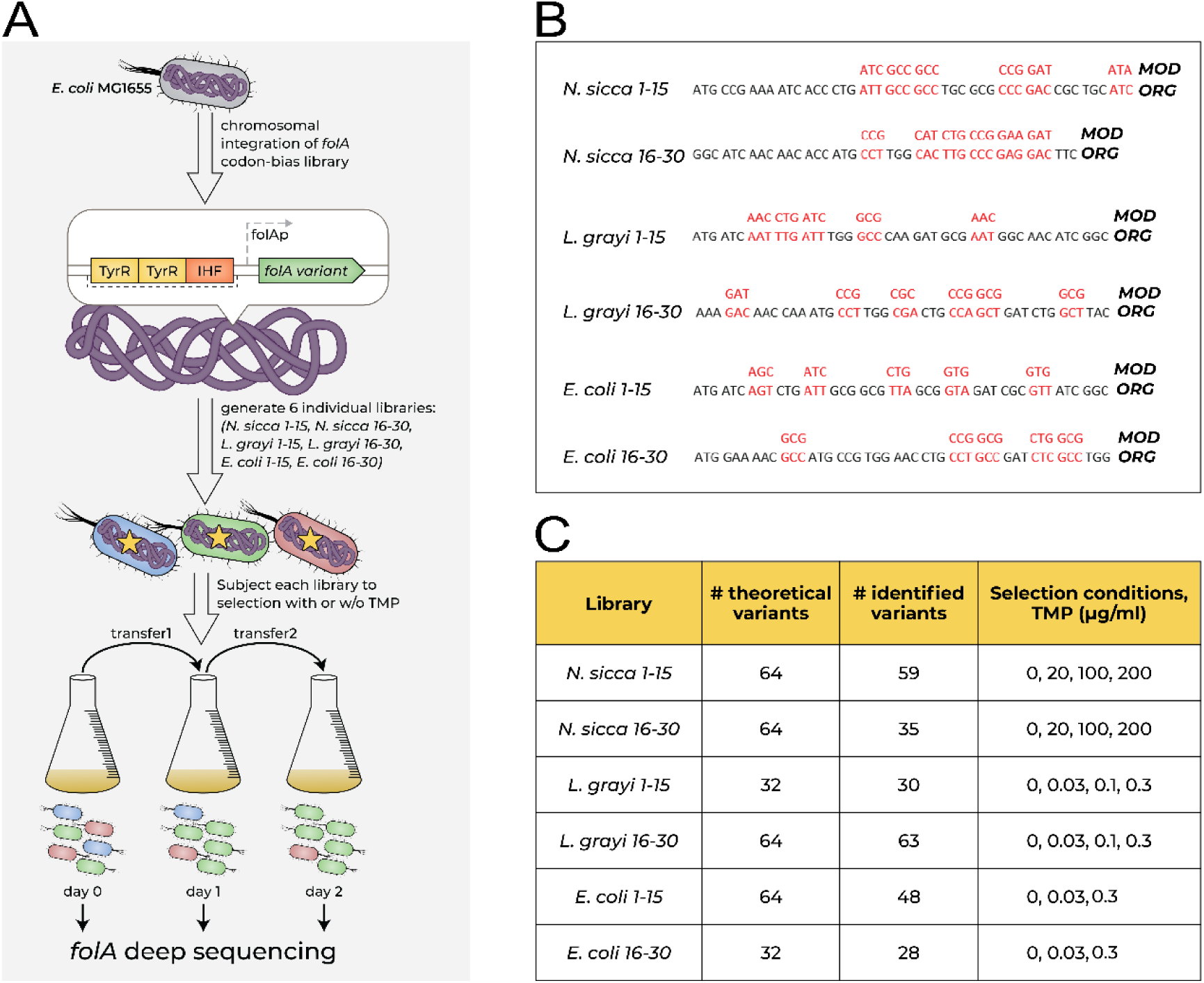
The competition experiment. (*A*) The experimental setup. (*B*) Codons diversified in each library (marked in red). (*C*) The theoretical number of expected codons, total number of the identified codons, and a range of TMP concentrations used in selection are shown for each library.

From changes in the relative frequencies of the individual library variants that occurred during the two consecutive passages (from day0 to day1, and from day1 to day2) (**table S4**), we calculated the distribution of fitness effects as well as the changes in the population average fitness and population Shannon diversity (see **Materials and Methods**) (**figs. 3,4** *upper panels*). Analysis of the distribution of the fitness effects within libraries subjected to selection reveals that variants carrying the full combination of the original codons within the diversified areas of *folA* genes were not necessarily the fittest (*i.e.*, reaching the highest frequency increase upon selection), or even more fit than the fully modified variants, which indicates that selection on synonymous codons of the transferred genes has the potential to further improve fitness of the host (**figs. 3,4** *lower panels*). Furthermore, in *L. grayi* and *N. sicca* libraries, selection with increasing amounts of TMP produced a gradual increase in the population average fitness and a decrease in the population diversity after only a single selection passage (from day0 to day1) (**figs. 3A,B**, **4A,B**). The gradual increase in selection stringency with the increasing TMP levels was apparent from the changes in the distribution of fitness effects of the individual library variants, a higher fraction of which became disadvantageous (*i.e.,* relative fitness drops below 1) at higher TMP levels (**figs. 3A,B**, **4A,B**).

**Fig. 3.**
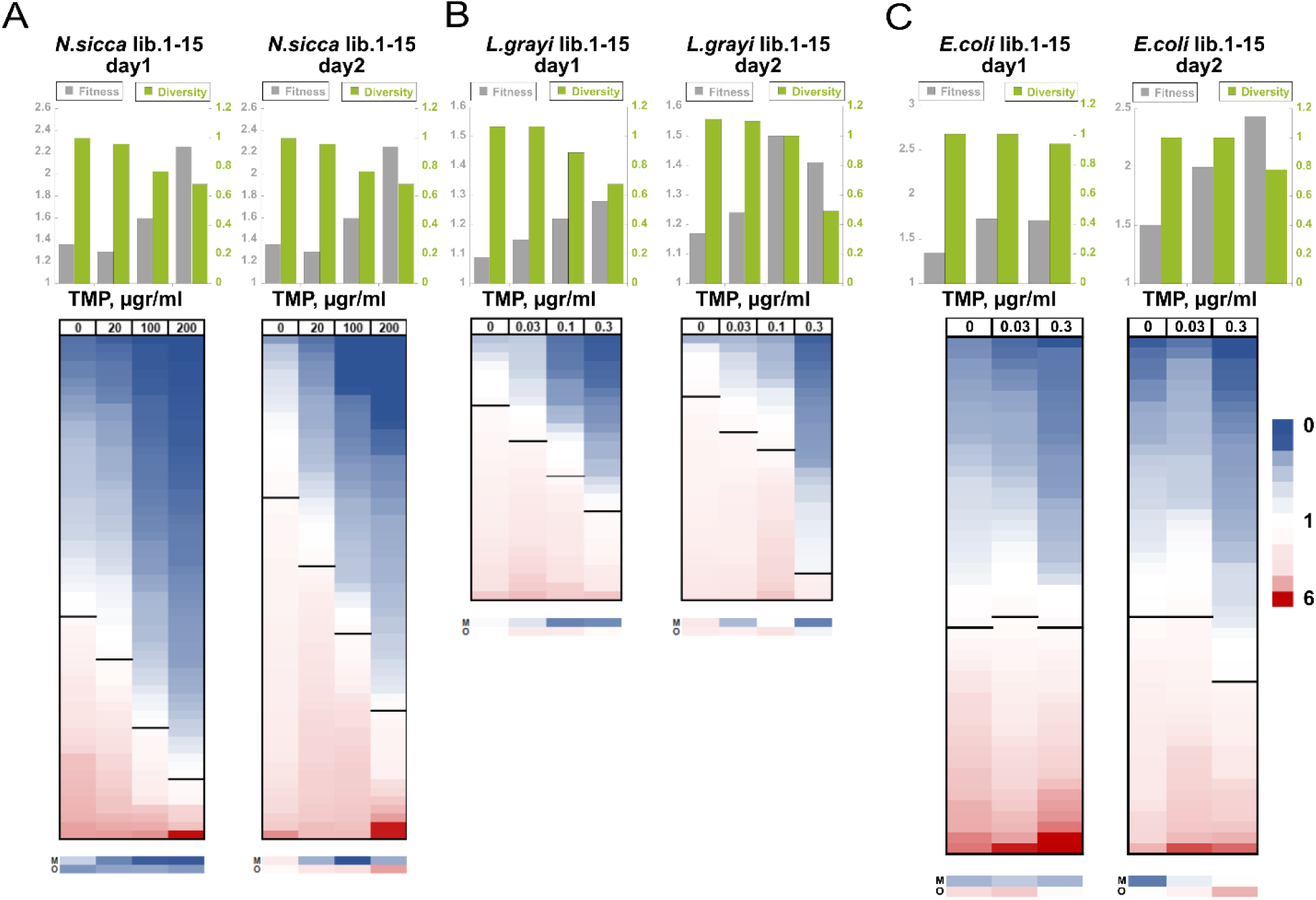
Codon bias drives population levels effects upon selection on *N. sicca*, *l. grayi*, and *E. coli* lib. 1-15. (*A-C*) *Upper panels*. Changes in the population relative fitness (*gray*) and Shannon diversity (*green*) after day1 and day2 of selection in the presence of 0, 20, 100, 200 μg/ml of TMP in case of *N. sicca*, 0, 0.03, 0.1 and 0.3 μg/ml of TMP in case of *L.grayi*, and 0, 0.03, and 0.3 μg/ml of TMP in case of *E. coli* libraries. (*A-C*) *Lower panels*. The relative fitness of individual library variants was sorted from lowest (*blue*) to highest (*red*) independently at each selection regime and presented as heat maps. (Note that variants at each row are not necessarily identical, as their fitness may change upon change in the selection regime). Black bars separate variants whose fitness dropped upon selection (fitness < 1) from those whose fitness has increased (fitness > 1). M, corresponds to variants with fully MOD sequence. O, corresponds to variants in which all synonymous codons encoding amino acids 1-15 are represented by ORG codons. The heat map ruler indicates the fold change in the relative fitness of the individual variants.

**Fig. 4.**
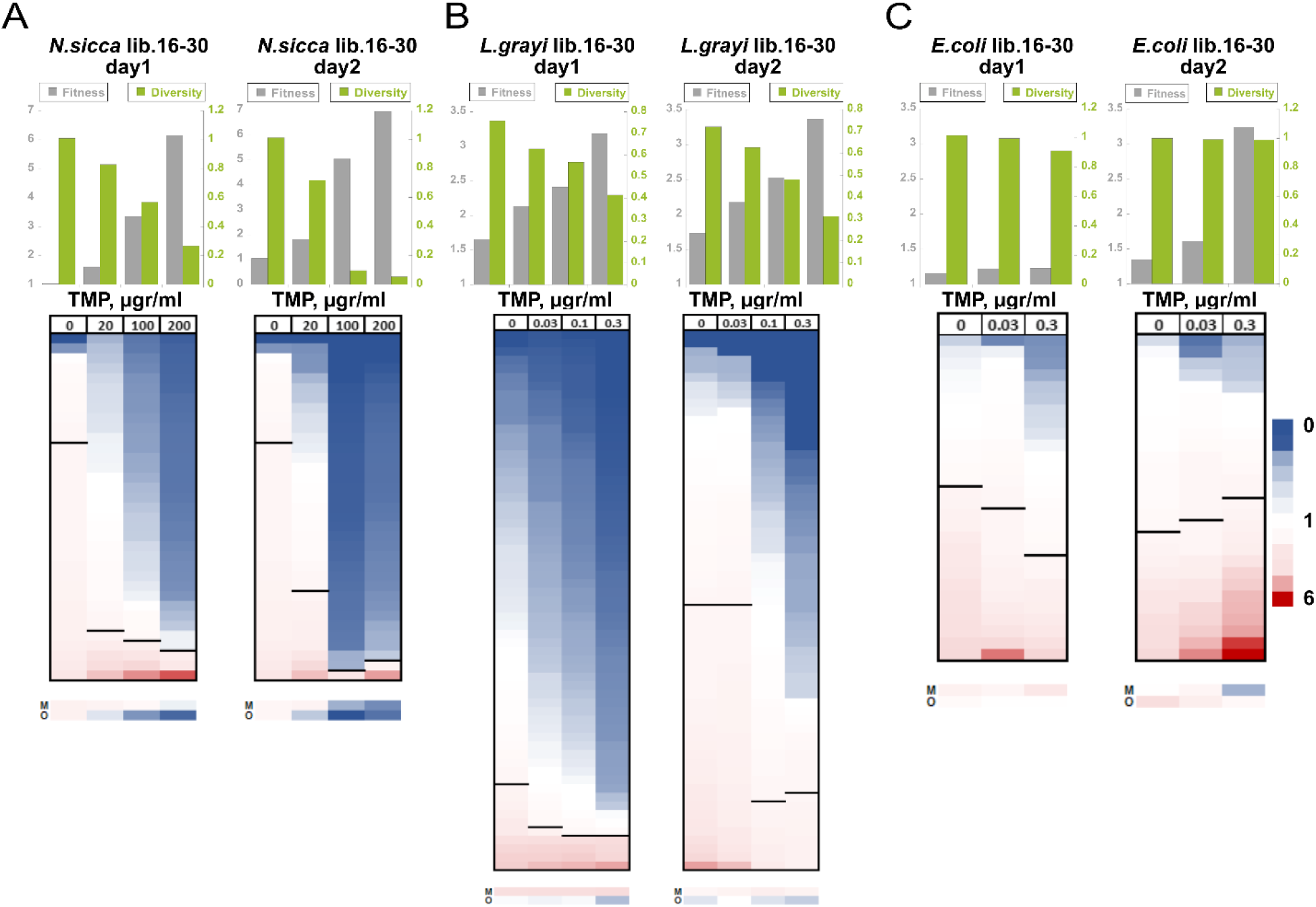
Codon bias drives population levels effects upon selection on *N. sicca*, *l. grayi*, and *E. coli* lib. 16-30. (*A-C*) *Upper panels*. Changes in the population relative fitness (*gray*) and Shannon diversity (*green*) after day1 and day2 of selection in the presence of 0, 20, 100, 200 μg/ml of TMP in case of *N. sicca*, 0, 0.03, 0.1 and 0.3 μg/ml of TMP in case of *L.grayi*, and 0, 0.03, and 0.3 μg/ml of TMP in case of *E. coli* libraries. (*A-C*) *Lower panels*. The relative fitness of individual library variants was sorted from lowest (*blue*) to highest (*red*) independently at each selection regime and presented as heat maps. (Note that variants at each row are not necessarily identical, as their fitness may change upon change in the selection regime). Black bars separate variants whose fitness dropped upon selection (fitness < 1) from those whose fitness has increased (fitness > 1). M, corresponds to variants with fully MOD sequence. O, correspond to variants in which all synonymous codons encoding amino acids 16-30 are represented by ORG codons. The heat map ruler indicates the fold change in the relative fitness of the individual variants.

Interestingly, the impact of selection was generally more pronounced in *L. grayi* and *N. sicca* libraries diversified within codons 16-30 than within codons 1-15. For instance, in *N. sicca* lib. 1-15, the average fitness has increased by 2.3-fold upon day0 to day1 passage in the presence of 200 μg/ml TMP versus 6.2-fold in lib. 16-30 (**figs. 3A, 4A**). Such differences can be explained by a stronger clonal interference between variants that comprise libraries 1-15. Clonal interference between multiple variants with comparable fitness effects delays the increase in the population average fitness by suppressing the selection sweeps by individual variants. Indeed, in lib. 1-15 of *N. sicca*, the relative frequency of the most fit variant in the library after a single passage selection in the presence of 200 μg/ml TMP increased to only 19.7% (from the initial frequency of 3.5% at day0). The relative frequency of this variant increased to 30% after the second passage (**table S4**). Conversely, under the same conditions, the most fit variant in lib. 16-30 reached a relative frequency of 85% on day1 (from initial frequency of 11.4% at day0). On day2, this variant almost fully swept the population, reaching over 98% in frequency (**table S4**). These dynamics are also captured by changes in the population diversity. The normalized Shannon diversity index of *N. sicca* lib. 1-15 (relative to the diversity of the naïve library that was set as 1) dropped to 0.63 after a single passage selection with 200 μg/ml TMP (**fig. 3A**). For comparison, the diversity of the *N. sicca* lib. 16-30 under the same conditions dropped to 0.22, nearly a 3-fold decrease (**fig. 4A**). In contrast to *N. sicca* and *L. grayi* libraries, *E. coli* libraries 1-15 and 16-30 did not respond to a single passage selection in the presence of increasing TMP concentrations in a gradual manner, showing only modest changes in the population average fitness and comparable distributions of fitness effects in the absence and presence of TMP (**figs. 3C, 4C**). Both *E. coli* libraries (1-15, and 16-30) also maintained a high Shannon diversity index, even after two selection passages, suggesting that they are subjected to a much higher impact of clonal interference than the xenologous libraries (**figs. 3C**, **4C**).

Since the shift in frequencies of competing variants was measured after a realtively short period of selection (the effect of selection became apparent in the xenologous libraries already after a single 16 hours-long passage), the calculated fitness effects could be, in theory, affected by the bias in the variant frequencies existing within the naïve libraries prior to selection (**fig. S9**). Thus, to estimate the role of the initial variant frequencies on the outcome of selection and to confirm the contribution of clonal interference to the observed evolutionary dynamics, we performed evolutionary simulations using SodaPop (Gauthier, et al. 2019) (see **Materials and Methods**). SodaPop is a forward-time simulator of the evolution of large asexual populations that entails a user-defined fitness landscape. The simulations were run using either the experimentally determined initial frequencies of *N. sicca* and *L. grayi* libraries 1-15 and 16-30, or, alternatively, with equalized initial frequencies. We found that the dynamics of changes in the average fitness and the Shannon diversity of the simulated populations largely recaptured the experimentally obtained dynamics (**fig. S9**). The simulation outcomes were found to be insensitive to the initial frequencies of the individual variants, thus validating the experimental results. Furthermore, the contribution of clonal interference to the observed dynamics of fitness changes in simulated populations was evident from the inverse linear correlation between the average fitness and the diversity of the evolving populations (**fig. S10**). At a higher selection stringency (higher TMP levels), the increase in the average population fitness was accompanied by a steeper decrease in population diversity, as expected under a clonal interference regime (**fig. S10**).

These findings show that in the xenologous *folA* libraries there exist two markedly different evolutionary dynamics, one for variants diversified within codons proximal to the translation start (codons 1-15) and another for variants diversified within codons 16-30. The difference in dynamics stems from the co-existence of multiple variants with comparable fitness effects in libraries 1-15, which delays selection sweeps by individual variants, as opposed to libraries 16-30 that are comprised of several variants with high fitness values that rapidly take over the populations. The clonal interference in libraries 1-15 also explains the gradual change in the evolutionary dynamics in response to an increase in TMP levels. The fitness effects of individual variants become more dissimilar with the elevation of stringency of selection. It is evident that in *E. coli folA* libraries, clonal interference is overall stronger, while being equally pronounced in both the 1-15 and 16-30 libraries.

### mRNA stability is the major determinant of codon bias-induced fitness effects upon HGT

Next, we determined whether the observed changes in fitness effects of the individual variants upon selection can be explained by their mRNA thermodynamic stability, CAIg/tAIg values, or GC content. We dismissed the possibility that the codon bias of the library variants affected fitness by controlling the co-translation folding of DHFR, since the diversified codons in our libraries cover only the first 30 amino acids. The diversified part of the nascent polypeptide chains in its entirety is therefore expected to fill the ribosome tunnel, known to accommodate 30-40 residues (Jacobs and Shakhnovich 2017). In all of the libraries diversified within codons 1-15 (*N. sicca* lib. 1-15, *L. grayi* lib. 1-15, and *E. coli* lib. 1-15), we established a strong correlation between the fitness of individual variants and their mRNA stability. mRNA stability was calculated using either a single 145 nt-long fragment covering the diversifications in codons 1-30 (from nucleotide −25, the beginning of transcription, and up to nucleotide +120), or via a 30 nucleotide-long sliding window starting from the mRNA transcription start site (nucleotide −25), in steps of 1 nt for a total of 110 windows (see **Materials and Methods**). Regardless of the method used for calculating mRNA stability, in all 3 libraries diversified within codons 1-15, selection unequivocally favored variants with less stable mRNA (**figs. 5A, S11A, S12A, S14A, S15A, S17A**). Furthermore, the correlation between the folding stability of mRNA transcripts and the fitness effects was highly dependent on TMP concentration. *E.g.*, in *N. sicca* up to 47% in fitness variance could be explained by mRNA stability at 200 μg/ml TMP upon day1 of selection (Spearman’s correlation *r* = 0.68; *p-value* = 2.54⸱10^-9^), whereas no correlation between mRNA stability and fitness was detected at 0 TMP (**fig. 5A**). Similar trends to those observed after day1 of selection were also detected when fitness effects were calculated for day2 (**figs. 5A, S13A, S16A**).

**Fig. 5.**
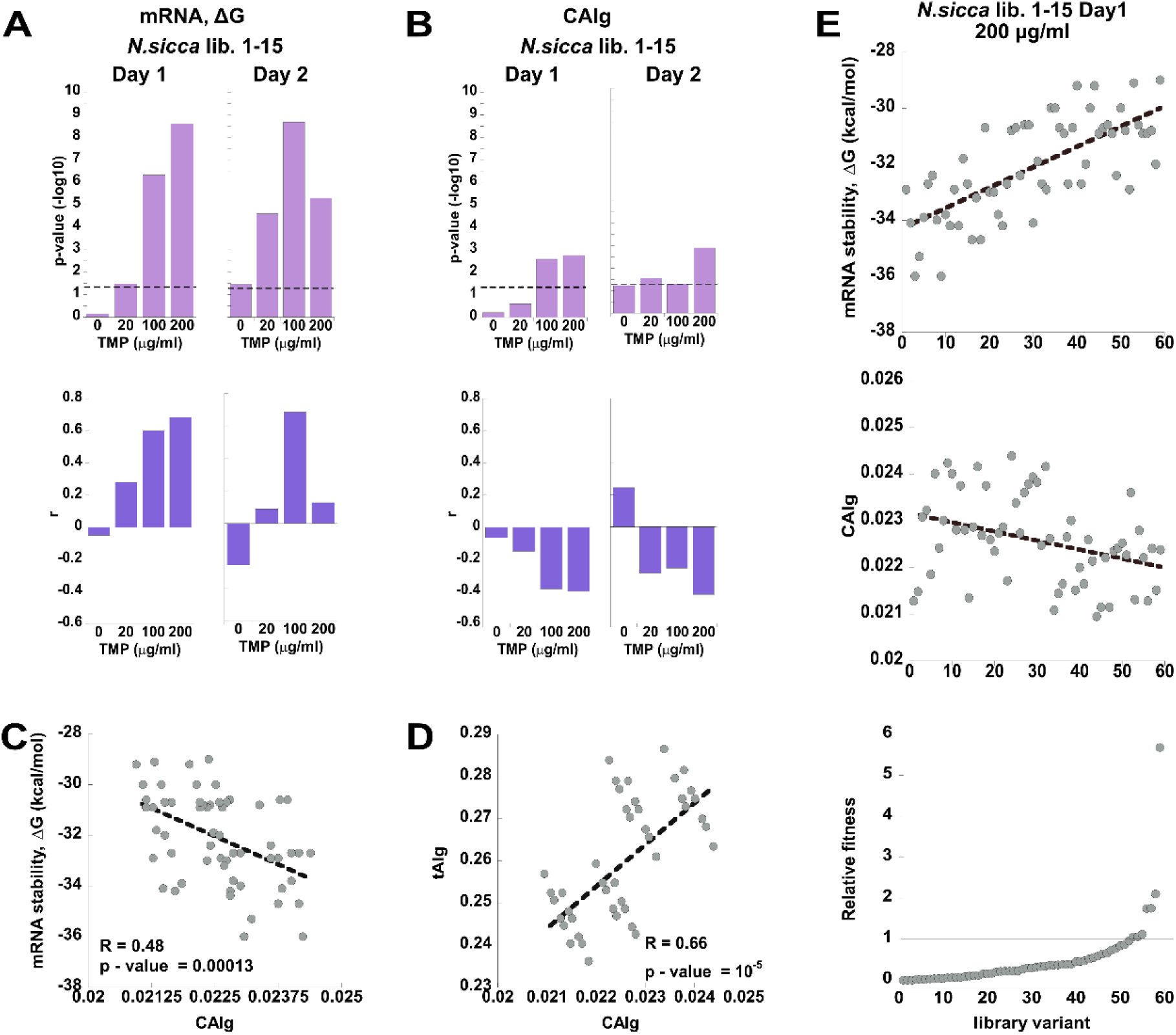
The selection on *N. sicca* lib. 1-15 is driven by 5’-mRNA stability. (*A,B*) *Upper panels. P-values* for Spearman non-parametric test for the association between the fitness effects and (*A*) the mRNA stability (calculated for a single fragment spanning 114 nucleotides from nucleotide −25, the beginning of transcription, and up to nucleotide +90), and the fitness effects and (*B*) the codon optimality (CAIg). *Lower panels.* Corresponding Spearman’s correlation coefficients, *r.* (*C,D*) Pearson’s correlations between (*C*) The mRNA stability and codon optimality (CAIg) and (*D*) between CAIg and tAIg values for variants comprising naïve libraries (day0). Pearson’s p-values and R values are indicated. (*E*) *Lower panel.* All library variants are sorted according to their relative fitness (from lowest to highest values) upon selection at 200 μg/ml TMP, day1. *Middle panel*. CAIg values of variants are shown according to their order in the lower panel. *Upper panel.* mRNA stability values of variants are shown according to their order in the lower panel.

In addition to mRNA stability, the changes in fitness effects in lib. 1-15 also correlated, albeit much less significantly, with CAIg values in *N. sicca* (Spearman’s correlation *r* = −0.4; *p-value* = 0.002; 200 μg/ml TMP, day1) (**fig. 5B**), with CAIg/tAIg values and GC content in *L. grayi* (Spearman’s correlation *r* = −0.42; *p-value* = 0.02; 0.3 μg/ml TMP, day1 for tAIg-fitness correlation) (**figs. S13A, S13A**), and with GC content in *E. coli* (Spearman’s correlation r = −l0.42; *p-value* = 0.03; 0.03 μg/ml TMP, day1) (**figs. S15A, S16A**). Which of these parameters, mRNA stability, codon optimality (expressed in either CAI or tAI scales), or GC content, then constitute the primary driver of fitness effects observed in libraries 1-15? To address this question, we first note that the correlation between fitness and CAIg/tAIg values is inverse, indicating that variants with more optimal codons are less fit, which contradicts the assumptions that codon optimality should produce more efficient translation, and, therefore, higher TMP resistance (**figs. 5B, S12A, S13A, S15A, S16A**). Second, we note that prior to selection (day0), there exists a significant inherent inverse correlation between mRNA stability and either CAIg or tAIg values among variants of the naïve libraries (**figs. 5C, S12C, S15C**), so that less stable mRNA variants have lower CAIg/tAIg values. The same inherent linkage between codon optimality and mRNA stability at gene starts was demonstrated for most bacterial strains with GC content higher than 50% (Bentele, et al. 2013), and is explained by the simple fact mRNA stability increases with increasing GC content. Consequently, selection to reduce secondary mRNA structure favors the incorporation of AU-rich codons. Such codons also happen to be rare (*i.e.*, less optimal) in bacteria with %GC >50. Thus, synonymous codons found in the vicinity of gene starts are sub-optimal primarily because they control translation by reducing the secondary structure of mRNA transcripts, and not because low codon optimality *per se* is advantageous at the starts of genes (Bentele, et al. 2013; Goodman, et al. 2013). This statement is supported by a strong inverse correlation between mRNA stability and GC content and/or by strong positive correlation between CAIg/tAIg values and GC content among naïve variants of of libraries 1-15 (**figs. S12C-E, S15C-E**). The similarity in correlation trends between the CAIg and tAIg values is also explained by the strong inherent correlation between them in most naïve libraries (**figs. 5D, S12F, S15F**). It is, therefore, plausible that the observed selection on codon bias in libraries 1-15 is driven primarily by the thermodynamic stability of mRNA transcripts, specifically by favoring variants with reduced secondary structure. The accompanying inverse correlation of fitness effects with codon optimality and GC content most probably also stems from the inherent correlation between stability, codon optimality, and GC content of the library variants prior to selection (**fig. 5E**). To validate that our conclusion is not a result of the bias in the variant frequencies in the naïve libraries, we introduced a bias of up to 10-fold in the initial frequency of variants in *N. sicca* lib. 1-15 (**fig. S18A**) and repeated the selection at 0 and 100 μg/ml TMP (**fig. S18B**). Similarly to results obtained with the unperturbed library, we found that the observed fitness effects in the presence of TMP were best explained by 5’-mRNA stability, with less stable variants being favored by selection (Spearman’s correlation *r* = 0.47; *p-value* = 0.62⸱10^-5^; 100 μg/ml TMP, day1) (**fig. S18C**). Furthermore, fitness effects also correlated with CAIg values, but as previously observed, the correlation was inverse and less significant than the one obtained with mRNA stability (Spearman’s correlation *r* = −0.32; *p-value* = 0.02; 100 μg/ml TMP, day1) (**fig. S18C**).

Analysis of the libraries diversified with codons 16-30 revealed that in *L. grayi* and *E. coli*, mRNA stability remains the most dominant force underlying the observed fitness effects (**figs. S12-17B**). However, in both libraries the significance of the mRNA stability – fitness correlation was substantially reduced in comparison to that of libraries 1-15. Depending on the day of selection, the correlation passed the significance threshold only when mRNA stability was measured using a 30 nt-long sliding window, and only in some of the selection conditions (**figs. S14B, S17B**). Similarly to dynamics observed in libraries 1-15, changes in the fitness effects in *L. grayi* and *E. coli* lib. 16-30 were inversely correlated with CAIg, tAIg and/or GC content on either day1 or day2 of selection (**figs. S13B, S14B, S16B, S17B**). As discussed above, this behavior can be explained by the inherent correlations that exist among the naïve library variants, namely those between mRNA stability and CAIg values (**figs. S12G, S15G**), mRNA stability and GC content (**figs. S12H, S15H**), CAIg and GC content (**figs. S12I, S15I**), and tAIg and CAIg values (**figs. S12J, S15J**).

Analysis of the *N. sicca* lib. 16-30 produced markedly opposite trend to that obtained with *L. grayi* and *E. coli* lib. 16-30. In *N. sicca*, no apparent correlation between mRNA destabilization and improved fitness could be observed neither on day1 nor on day2 of selection (**figs. 6A**). In fact, when mRNA folding stability was calculated using a 30-nt sliding window, more stable variants appeared to be weakly favored by selection (windows #40-50 on day1, and windows #70-80 on day2, **fig. S11B**). In addition, fitness effects obtained upon selection at 20 μg/ml TMP on day1 and at 100 and/or 200 μg/ml TMP on day2 significantly correlated with codon optimality. As evidenced by tAIg and/or CAIg metrics, or by GC content, variants that carry more optimal codons or those that exhibit a higher GC content were found more fit (Spearman’s correlation *r* = 0.25, *p-value* = 0.009 for tAIg-fitness correlation; *r* = 0.55 *p-value* = 0.0007 for GC content-fitness correlation on day1 and *r* = 0.5, *p-value* = 0.003 for CAIg-fitness correlation; *r* = 0.05, *p-value* = 0.002 for tAIg-fitness correlation; *r* = 0.45 *p-value* = 0.006 for GC content-fitness correlation on day2) (**figs. 6B**, **S19A,B**). The positive correlation between codon optimality and fitness effects directly supports the notion that codon optimality controls translation efficiency and, therefore, is favored by selection. The similarity between correlation trends obtained with CAIg and tAIg values is not surprising, given that both metrics are highly correlated among the naïve variants (**fig. 6D**). Notably, no inherent correlation is observed between mRNA stability and codon optimality in variants that make up the naïve library (**fig. 6C**). It, therefore, appears that in *N. sicca* lib. 16-30, selection towards codon optimality was the driving force behind the observed changes in fitness effects (**fig. 6E**). The correlation between fitness effects and GC content is also most likely driven by codon optimality due to the significant positive correlation observed between the tAIg values and GC content among the naive variants comprising the library (**fig. S19C**).

**Fig. 6.**
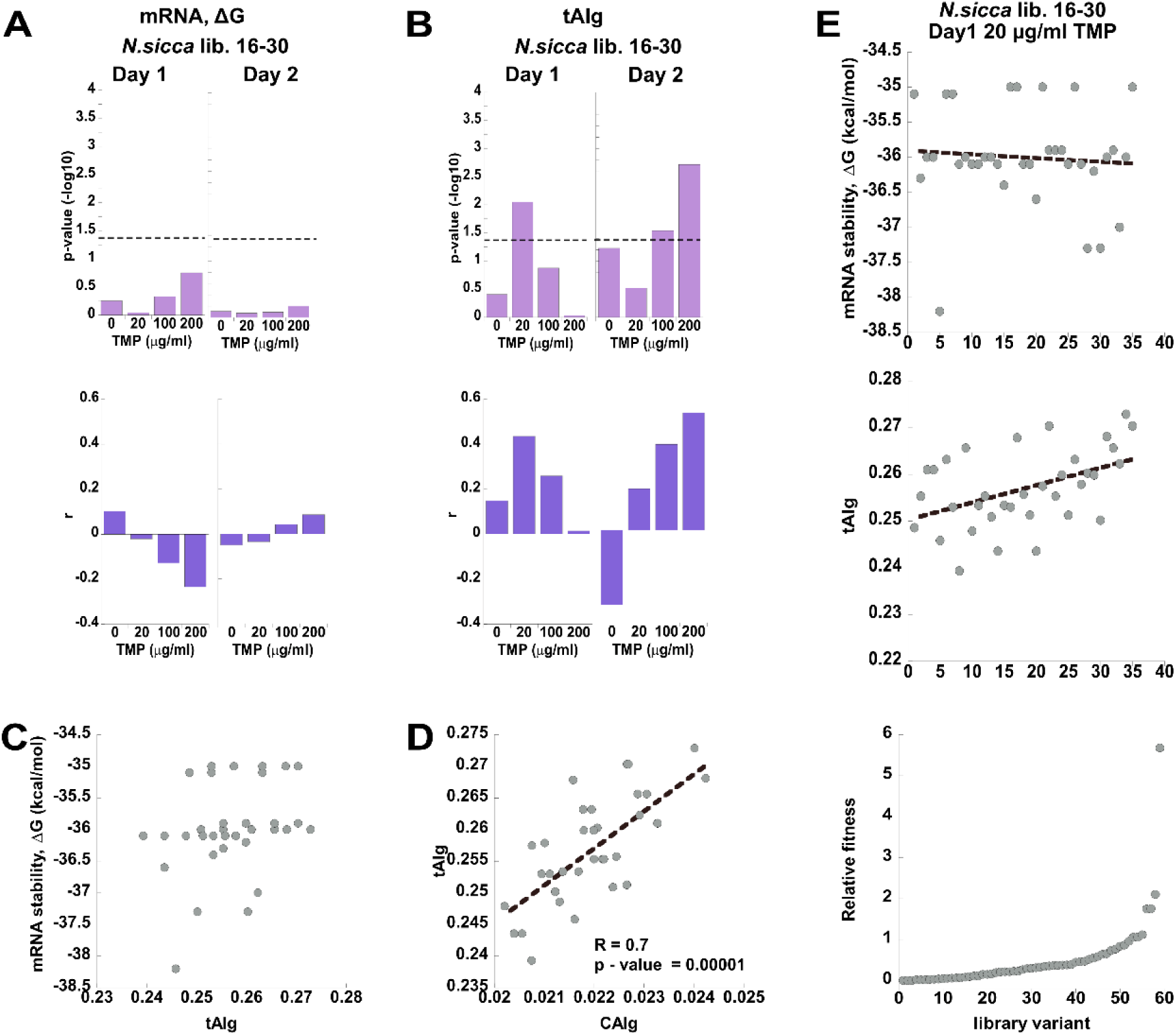
The selection on *N. sicca* lib. 16-30 is driven by codon optimality. (*A,B*) *Upper panels. P-values* for Spearman non-parametric test for association between fitness effects and (*A*) mRNA stability (calculated for a single fragment spanning 114 nucleotides from nucleotide −25, the beginning of transcription, and up to nucleotide +90), and fitness effects and (*B*) codon optimality (tAIg). *Lower panels.* Corresponding Spearman’s correlation coefficients, *r.* (*C,D*) Correlations between (*C*) mRNA stability and codon optimality (CAIg) and (*D*) between CAIg and tAIg values for variants comprising naïve libraries (day0). Pearson’s p-values and R values are indicated only for the significant correlation (p < 0.05). (*E*) *Lower panel.* All library variants are sorted according to their relative fitness (from lowest to highest values) upon selection at 20 μg/ml TMP, day1. *Middle panel*. tAIg values of variants are shown according to their order in the lower panel. *Upper panel.* mRNA stability values of variants are shown according to their order in the lower panel.

Collectively, the presented findings suggest that both mRNA stability and codon optimality contribute to the compatibility of the transferred *folA* genes. However, their impact is highly dependent on the environmental conditions, *i.e.*, the concentration of TMP. Although *N. sicca* and *L. grayi* DHFR differ substantially in TMP sensitivity, there exists a range of sub-MIC TMP concentrations for each xenologous strain whereby codon bias-related fitness effects may be predicted directly from sequence changes. mRNA stability dominates the codon bias-related cause of fitness changes. When HGT leads to 5’-end overstabilization of mRNA transcripts, mRNA stability outweighs the fitness contribution of codon optimality.

### 5’-end overstabilization of mRNA transcripts can cause mRNA accumulation outside the polysome

Having established the dominant role of 5’-end mRNA folding stability in codon bias compatibility of the transferred genes, we were interested to see whether changes in mRNA stability can affect mRNA abundance. To this end, we measured the changes in the steady-state mRNA abundance upon synonymous codon replacement in all *folA* xenologous strains (**fig. 7A**). In the absence of TMP, the effect of codon replacement on mRNA levels was highly variable. Some xenologues genes exhibited a severe to mild decrease in mRNA levels, if any change was observed at all (*N. sicca*, *M. capsulatus*, *P. putida, M. mesenteroides*). Other xenologues exhibited an increase in mRNA levels (*B. subtilis, L. lactis, L. grayi*). The addition of TMP had no effect on this trend (**fig. S20A**). Surprisingly, the increase in the steady-state levels of mRNA transcripts upon codon replacement was not necessarily matched by a corresponding increase in protein levels (**fig. 1B**). For instance, app. 5-fold increase in mRNA abundance of *folA* from *L. lactis* and *L. grayi* was accompanied by a 4-6-fold decrease in the protein levels (**figs. 1B, 7A**).

**Fig. 7.**
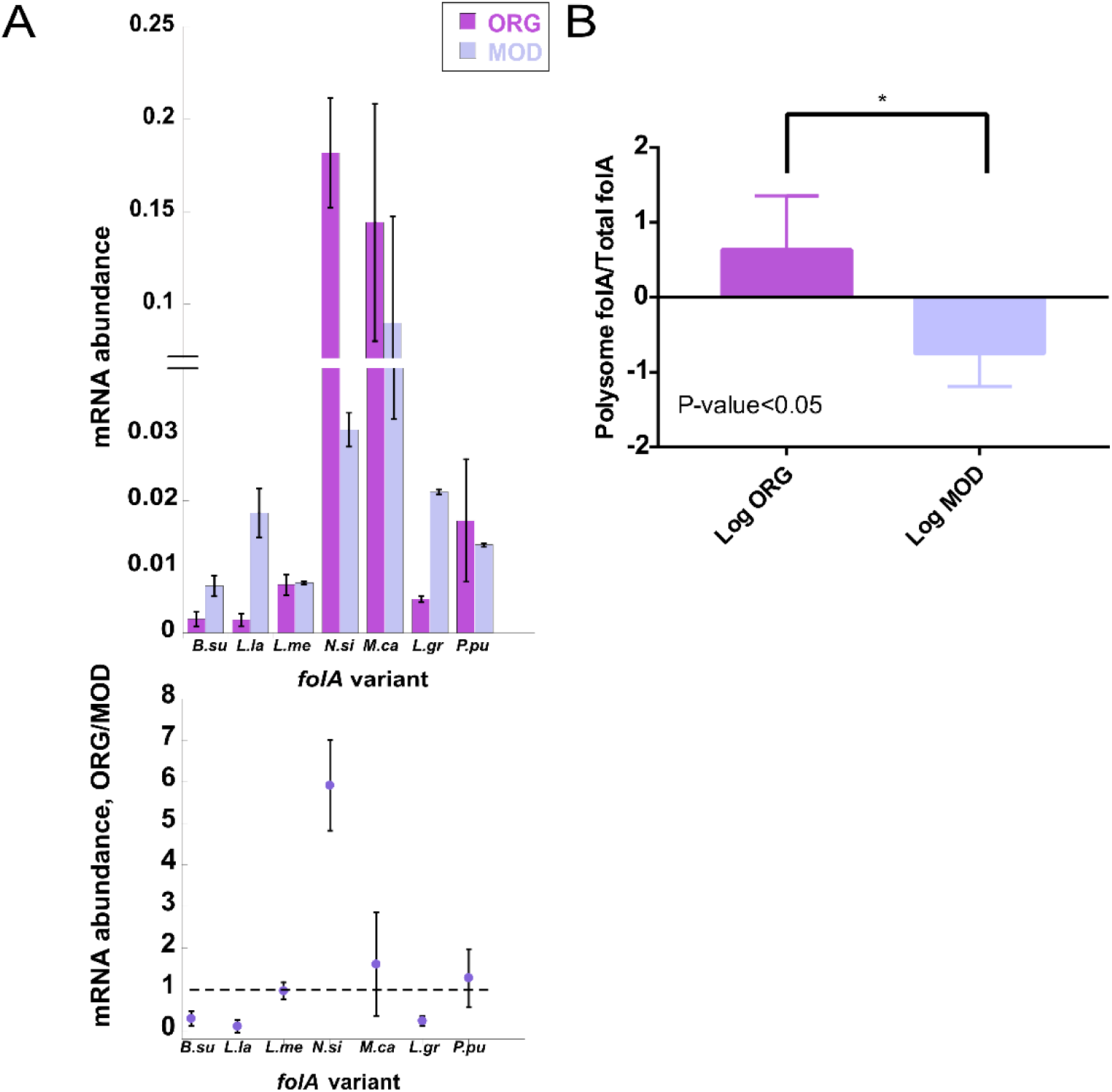
The effect of codon usage of *folA* xenologoues on mRNA abundance in *E. coli* host. *folA* variants are marked as in **table 1**. (*A*) *Upper panel.* Intracellular steady-state levels of original (ORG) and modified (MOD) *folA* mRNA. *Lower panel*. Ratios of *folA* mRNA abundance between ORG and MOD *folA* sequences. (*B*) Log ratios between *L. grayi folA* mRNA abundance found within the polysomal and total fractions calculated for ORG and MOD sequences. Error bars were calculated from four independent measurements.

How can such an inconsistency be explained? Since the inefficient ribosome translation elongation triggered by codon usage-incompatible mRNA is expected to cause transcript decay (Presnyak, et al. 2015), most plausibly through the involvement of Rho factor (Richardson 1991), we hypothesized that the lack of mRNA decay observed on the background of inefficient translation is a result of the reduced engagement of mRNA transcripts with translating ribosomes. In other words, mRNA transcripts escape degradation and accumulate due to the reduced participation in translation. Thermodynamic overstabilization of the endogenous *folA* mRNA structure near the translation start codon was indeed shown to impede translation initiation by hindering the accessibility to Shine-Dalgarno binding site in *E. coli* (Bhattacharyya, et al. 2018). Although in this example the mRNA transcripts unengaged in translation were efficiently degraded, it is plausible that the overstabilization of the 5’-end of some of the xenologous *folA* transcripts not only limits the accessibility of the ribosomal binding site but also conceals potential cleavage sites recognized by endonucleases, thus reducing the degradation propensity of the transcript (Rauhut and Klug 1999). To test this hypothesis, we extracted the total and polysomal mRNA fractions from mid-exponentially growing MOD and ORG *L. grayi* strains and quantified the ratios of the polysomal and the total *folA* mRNA steady-state levels in both strains (see **fig. S20B** and **Materials and Methods**). As hypothesized, we observed a significant shift of the MOD *L. gayi folA* mRNA transcripts from the polysomal to the total fraction, indicating that the accumulated transcripts indeed avoid translation (**fig. 7B**). The aggregation resistance of the MOD transcript may result from the overstabilization of its 5’-end by 5 kcal/mol (**fig. S5F**).

Our findings suggest that mRNA decay is not solely the immutable outcome of the codon bias-induced reduction in translation efficiency of the horizontally transferred genes, as was previously thought. The inefficient translation and transcript accumulation can occur simultaneously, resulting from the poor engagement of transcripts by translating ribosomes. Thus, steady-state mRNA abundance cannot serve as a reliable predictor of translation efficiency in the case of HGT.

## Discussion

To disentangle the contribution of codon bias to the functional integration of horizontally transferred genes in bacteria from other barriers and constrains that control the outcome of HGT events, we designed an experimental system that establishes a direct link between the variability in codon usage of the transferred genes and fitness of the host. Using *E. coli* as a host, and the *folA* gene as a target for HGT via chromosomal xenologous replacements, we determined that mRNA stability at gene starts, and codon optimality have a profound effect on the fitness of the host. The fitness effects of codon bias within codons 1-15 were dominated by mRNA stability – selection strongly favored destabilized mRNA variants. Since mRNA destabilization is achieved by codons with reduced GC content, and because such codons also happen to be rare in bacteria with GC content >50%, selection of variants with less stable 5’-end mRNA structures was accompanied by an apparent enrichment of variants composed of less optimal codons. Importantly, in *N. sicca* and *L. grayi* libraries 1-15, the selection of less stable variants was acutely dependent on the environmental conditions, *i.e.,* levels of TMP in the growth media. Indeed, in both libraries, no enrichment of less stable mRNA was apparent in the absence of TMP. The selection intensified, however, following only one day of exposure to sub-lethal levels of TMP. Since *N. sicca* and *L. grayi* DHFRs differ in TMP sensitivity, the range of TMP concentrations that triggered selection for less stable mRNA variants did not overlap between the libraries (0.03 - 0.3 μg/ml TMP in case of *L. grayi* versus 20 – 200 μg/ml TMP in case of *N. sicca*).

The dominating role of mRNA stability was challenged by codon optimality in libraries diversified within codons 16-30. Whereas in *L. grayi* lib. 16-30 mRNA stability remained the main cause of codon bias fitness effects, in *N. sicca* lib. 16-30 the variability in fitness effects was best explained by the contribution of optimal codons, defined either as codon adaptation index (CAI) or tRNA adaptation index (tAI), with variants composed of optimal codons being favored by selection. As with mRNA stability, the contribution of codon optimality to codon bias-related fitness effects was apparent only within a relatively narrow range of TMP concentrations (**fig. 6A,B**). The change in TMP sensitivity upon codon replacement from original to frequent codons performed throughout the entire sequence of *folA* xenologoues also supports the dominating role of the 5’-end of mRNA stability over codon optimality. Indeed, although codon replacement led to an increase in codon optimality (either CAIg or tAIg values) in all seven xenologous strains (**fig. S4A,B**), only in the *E. coli* strain carrying MOD *B. subtilis* such a replacement was accompanied by an increase in resistance to TMP (**fig. S7A** and **table 2**). This xenologue also happened to be the only one that exhibited a substantial destabilization (around 5 kcal/mol) of its 5’- end mRNA structure upon codon replacement (**fig. S5A**). Thus, unless 5’-end mRNA is destabilized, the contribution of codon optimality to fitness in the xenologous strains is subdued. Quite intriguingly, we also found that 5’-end mRNA overstabilization of the transferred genes in some instances can lead to an accumulation of mRNA transcripts outside of the polysome, suggesting that codon bias-induced reduction in translation efficiency is not always accompanied by mRNA decay. Thus, mRNA abundance cannot serve a reliable proxy to translation efficiency of foreign mRNA transcripts.

Our findings suggest that although selection operates universally to reduce the thermodynamic stability at gene starts in all donor organisms, horizontal gene transfer may nonetheless result in the overstabilization at the beginning of mRNA transcripts and, consequently, in reduced production of foreign proteins within the host bacteria. In this scenario, the mRNA stability, which controls translation initiation, is the most dominant codon bias-related effect in HGT, superseding the impact of codon optimality on controlling translation elongation. This conclusion supports the idea that translation initiation constitutes the rate limiting step of protein synthesis by the ribosome.

## Materials and methods

### Chromosomal replacements of the *folA* xenologues

The chosen xenologous *folA* gene sequences fused to a tag encoding six histidines (**table 1** and **table S1**) were synthesized (Integrated DNA Technologies), cloned into vector pKD13 specifically designed for λ Red homologous recombination of *folA* genes (Bershtein, et al. 2012), and integrated into *E. coli* chromosome by replacing the endogenous *folA* coding sequence but preserving the upstream endogenous regulatory region. Recombinant strains were selected on LB agar plates supplemented with 25 μg/ml chloramphenicol and 35 μg/ml kanamycin and chromosomal integration was validated by Sanger sequences, essentially as described (Bershtein, et al. 2012).

### Media and growth conditions

All variants were grown overnight from a single colony in 5 ml M9 minimal media salts supplemented with 0.2% glucose, 0.1% casamino acids, 1mM MgS0_4_, 0.5 µg/ml thiamine at 37^0^C. Bacterial cultures were then diluted 1/100 and grown at 37^0^C for 12 hours in 96-well microtiter plates (16 wells for each strain) in the absence or presence of a range of concentrations of trimethoprim (Sigma). 600 nm OD data were collected at 10 min intervals. The resulting growth curves were fit to a bacterial growth model to obtain growth rates parameters (Zwietering, et al. 1990).

### IC_50_ determination

Growth was quantified by integration of the area under the growth curve (OD vs. time) between 0 and 15 h, as described in ref. (Rodrigues, et al. 2016). Growth integrals determined for a given variant were normalized in respect to the corresponding growth of that mutant measured in the absence of TMP. IC_50_ values were determined from the fit of a logistic equation to plots of growth vs. TMP concentrations.

### Intracellular protein abundance measurements

All variants were grown overnight from a single colony in 5 ml supplemented M9 minimal media at 37^0^C. Bacterial cultures were then diluted 1/100 and grown at 37^0^C for about 4 hours until the cells reached OD∼ 0.5. Cells were lysed with BugBuster (Novagen) and the total protein concentration within soluble fractions was determined by BCA (CYANAGEN). Lysates were then normalized by total protein abundance, loaded on Ni-NTA 0.2ml column (G-BIOSCIENCES), and DHFR proteins were isolated using manufacturer’s instructions. The intracellular DHFR amounts within soluble fractions were eventually determined by SDS-PAGE followed by standard Western Blot protocol using mouse anti-His monoclonal antibodies (abm) and goat anti-mouse HRP antibodies (abm).

### Intracellular mRNA abundance measurements

All variants were grown overnight from a single colony in 5 ml supplemented M9 minimal media at 37^0^C. Bacterial cultures were then diluted 1/100 and grown at 37^0^C for about 4 hours, until cells reached OD∼ 0.6. 1.5 ml of the cells were treated with RNAprotect Bacteria Reagent (QIAGEN). Total RNA was extracted using GeneJET RNA purification kit (Thermo scientific), flowed by DNAse I digestion (Thermo scientific). Total RNA concentration was determined by nanodrop (DeNovix). 400 ng of the purified RNA was subjected to iScript cDNA synthesis (BIO-RAD). Abundance of *folA* mRNA was estimated through quantitative PCR using SYBR green kit (KAPABIOSYSTEMS) (**table S5**, primers# 1-2). 16S rDNA gene was used as an internal reference (**table S5**, primers# 3-4).

### Promoter activity measurements

*MG1655* strains were transformed with pUA-66 plasmid containing *folA* promoter fused to green fluorescent protein-coding sequence (Zaslaver, et al. 2006). The strains were grown overnight in 5 mL M9 minimal media supplemented with 0.2% Glucose, 1 mM MgS0_4_, 0.5 µg/mL thiamine, 0.1% casamino acids and, diluted 1/100 and grown at 37^0^C for 12 hours in 96-well microtiter plates (16 wells for each strain) in the absence or presence of IC50 levels of trimethoprim (see **table 2**). 600 nm OD and GFP fluorescent signal (excitation 485 nm and emission 517 nm) data were collected at 10 min intervals. The ration between the fluorescent signal and biomass production (OD) was defined as promoter activity.

### mRNA folding stability calculation

mRNA folding stability (Gibbs free energy difference between folded and folded state, ΔG, kcal/mol) was calculated for either a single fragment spanning 145 nucleotides (from nucleotide −25, the beginning of transcription, and up to nucleotide +120) to cover the diversified codons 1-30, or using a 30 nucleotide-long sliding window (to reflect the size of a ribosome footprint) starting from the mRNA transcription start (nucleotide −25) in steps of 1 nt. mRNA stability calculations were performed with unafold software (Markham and Zuker 2008).

### Generation of libraries with codon diversification

Fully modified (MOD) *folA* sequences from *E.coli*, *L.grayi* and *N.sicca* were diversified back to the original codons (ORG) within codons 1-15 or 16-30 in combinatorial manner in positions indicated in **Fig. 2B**. The diversification was carried out by PCR performed with a mixture of primers diversified in the designated positions (primers #5-15, **table S5**). The products of PCR for each library were then amplified by primers # 17-18 (**table S5**), cloned into vector pKD13 specifically designed for λ Red homologous recombination of *folA* genes (Bershtein, et al. 2012), and integrated into *E. coli* chromosome by replacing the endogenous *folA* coding sequence while preserving the upstream endogenous regulatory region, essentially as described (Bershtein, et al. 2012). Integrants for each library were selected on LB agar plates supplemented with 25 μg/ml chloramphenicol and 35 μg/ml kanamycin. In total, between 685 to 2800 individual colonies were obtained for each library (**table S6**). The colonies were collected from the LB agar plates by scraping in the presence of 10 ml LB and cryopreserved at −80°C in the presence of 50% glycerol.

### Laboratory evolution experiment

The evolution experiment was performed for 16 hours for two consecutive days at a range of TMP concentrations (**fig. 2C**). The cryopreserved libraries (designated as day0) were defrosted, diluted 1/100 in 50 ml of supplemented M9 media, and grown for 16 hours at 37°C, 220RPM in a 250ml Erlenmeyer flask (designated as day1 of selection). This was followed by another 1/100 dilution in a fresh 50 ml of supplemented M9 media and growth for additional 16 hours at 37°C, 220RPM in a 250ml Erlenmeyer flask (designated as day2 of selection). Aliquots from each each time point were collected and cryopreserved at −80°C with addition of 50% glycerol. The frequencies of each variant in the libraries were then analyzed by deep sequencing in three time points (day 0 - initial frequencies, day1- after 16 hours of evolution, and day 2- after 32 hours of evolution).

### Deep sequencing

Sample preparation for the deep sequencing had three steps: first, *folA* gene from the evolved and naïve populations was amplified by PCR. 10µl of each aliquot were diluted in 100 µl ddw, and 1µl of the template was amplified in 50 μl PCR reactions supplemented with 4µl 2.5mM dNTP’s, 1µl (250U) PrimeStarGXL (TAKARA), and 5 μl of 10 μmol/μl “FWD 2 Deep seq” and “Rev Deep seq” primers (**table S5**). Second, the PCR products were run and purified from 1% agarose gel (Machary-Nagel) and used as template for the next amplification reactions performed in 50 μl with 5µl i5 and i7 from Nextera XT DNA library preparation kit (Illumina) (see **table S7** for Nextera primers used), 1µl (250U) PrimeSTAR GXL DNA polymerase (TAKARA), 4µl 2.5mM dNTP’s, and 55 ng DNA template DNA. The PCR program was: 94°C for 5 minutes (one cycle), followed by 12 cycles of 95°C for 10 sec, 55°C for 15 Sec and 68°C for 45 sec, and a single cycle of 68°C for 5 minutes. The products of the PCR reaction with Agencourt AMPure XP PCR purification kit (Beckman Coulter). The deep sequencing was performed by Nextera Miseq reagent micro kit v2 by MiSeq platform (Illumina). Raw reads (FASTQ files) were trimmed using fastp tool (Chen, et al. 2018). Next, any sequences in the mutagenized library that were shorter than the expected length were discarded. The generated fastq files were then analyzed with Enrich2 software (Rubin, et al. 2017) to determine the read count for each variant using the “count only” scoring option and the “wild type” normalization parameter.

### Population analysis

The frequency (f) of each variant in a specific time point was calculated by the division of the number of reads (n (t)) by the total number of read at this time point 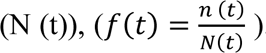. The fitness in each time point (w(t)) was calculated by the ratio between the frequencies of a variant in a specific time point (f (s)) and a reference time point 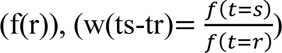. The relative fitness of each variant in a specific time point was calculated (w’ (ts-tr)) by multiplying the fitness (w (ts- tr)) and the frequency in the specific time point (f(r)), (w’ (ts-tr) =𝑤(𝑡𝑠 − 𝑡𝑟) ∗ 𝑓(𝑟)). The average fitness of the population in a specific time point was calculated as the sum of the relative fitness for all the variants in the populations for a specific time point 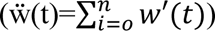 and it is normalized by adding the (ẅ_norm_(t-1)) value of the previous day ẅ and subtracting 1. The value of ẅ at the day 0 is defined as 1. The population Shannon diversity (H) was calculated as 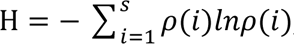, where *p* is the frequency of an individual variant *i*, and *s* is the number of variants.

### Statistics

The correlation between bacterial fitness and physical mechanisms that influence by synonymous mutations was calculated by spearman correlation and multiple linear regression. Spearman test was done between the fitness and geometric mean of CAI, tAI, mRNA stability and GC content at each time point for each population. The geometric mean of each population was calculated as the sum of geometric mean of each variant multiplied by the frequency of the variant in each time point (FigureS5). The output was rho and P-value. To determine the correlation between the local mRNA stability and bacterial growth spearman rank test was done between the fitness of the variants and each part of the mRNA sliding window. The output was rho and p-value for each part of the sliding window (R script is in the supplementary).

### Polysome extraction

Polysome extraction was done essentially as described (Liang, et al. 2018; Alasad, et al. 2020). *E. coli* strains carrying ORG and MOD *L. grayi folA* genes were grown overnight from a single colony in 5 ml supplemented M9 media at 37^0^C. Bacterial cultures were then diluted 1/100 in 50 ml of M9 media (in a in 250 ml Erlenmeyer flask) and grown at 37^0^C for about 4 hours until the cells reached OD∼ 0.6. The cultures were then poured into 50ml flasks (25ml to each flask) packed loosely with crashed ice. Cultures were quickly mixed with the ice by inverting the flask. Next, the cells were pelleted down by centrifugation (8000 RCF, 20 minutes, 4^0^C) and resuspended in 0.5 ml lysis buffer (10mM Tris-HCl pH8.0, 10mM MgCl2, 1mg/ml lysozyme 1mg/ml). The resuspended cells were flash frozen in liquid nitrogen and placed in an ice cooled water bath with occasional mixing. Once the cells melted, they were frozen again in liquid nitrogen and kept at −80^0^C for the next step. The cells were defrosted on ice, mixed with 15 µl sodium deoxycholate, and spanned down in pre-cooled microcentrifuge (10000RCF, 10 minutes, 4^0^C). The clarified portion was transferred to a fresh tube and mixed with 3 µl of RNaseOUT(Invitrogen). 50 µl were then collected, supplemented with 450µl DDW and 500 µl Trizol (Sigma), and stored at −80^0^C as the total mRNA fraction. The rest of the clarified portion was loaded on a sucrose gradient and subjected to ultracentrifugation (229884 xg, 2.5 hours, 4°C). The polysome profile was read using a piston gradient collector (Biocomp) fitted with a UV detector (Tirax). Three polysomal fractions were collected for each growth condition, placed in Trizol (Sigma), and stored at −80^0^C as the polysomal mRNA fraction. The RNA from the total and polysomal RNA fractions were extracted using manufacturer’s instruction and analysed by qPCR with primers for *folA* gene (rimers# 1-2, table S5) and *adk* gene (primers# 21-22, table S5), that was used as an internal control. The ratio between the threshold cycle of *folA* and *adk* was calculated (ΔCT = CT(folA)−CT(adk)) and was used to determine the relative abundance in each fraction 2^-2ΔCt^ . Increased association of the *folA* gene with polysomes will increase this value. Because of the noisy nature of Polysomal extraction method log2 transformation was applied on the results. log2 transformation of 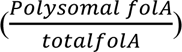 ratio was applied.

### Simulations

Evolutionary simulations were performed with SodaPop software (Gauthier, et al. 2019). The simulations started with an initial size of 10^6^ cells, partitioned into the library variants using either equalized or experimentally determined variant frequencies at t=0. Then, the frequency of each variant was updated using a standard Moran process (Moran 1958), where the fitness of each cell quantitatively represents the doubling time of each cell. This fitness represents a ‘fitness score’ estimated using 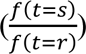, starting from the experimental read counts. To mimic the fluctuation in experimental population sizes due to daily dilutions upon passaging, we let the simulated populations to reach 10^8^ cells over 12 generations, followed by dilution 1/100 dilution every six generations. The dilution is implemented as a random down sampling. Similar to the experiment, in this short number of generations, we assumed that no *de novo* mutations occurred in the library and that all variant frequency changes are either due to its initial fitness or neutral drift (fully implemented in the explicit Moran process by SodaPop). We performed ten replicate simulations for each of the four experimental conditions corresponding to TMP concentrations.

## Acknowledgements

We thank Adrian I. Levy for his feedback on the manuscript. S.B. was supported by the Israel Science Foundation personal research grant 593/21.

## Supplementary Figures

**Fig S1.**
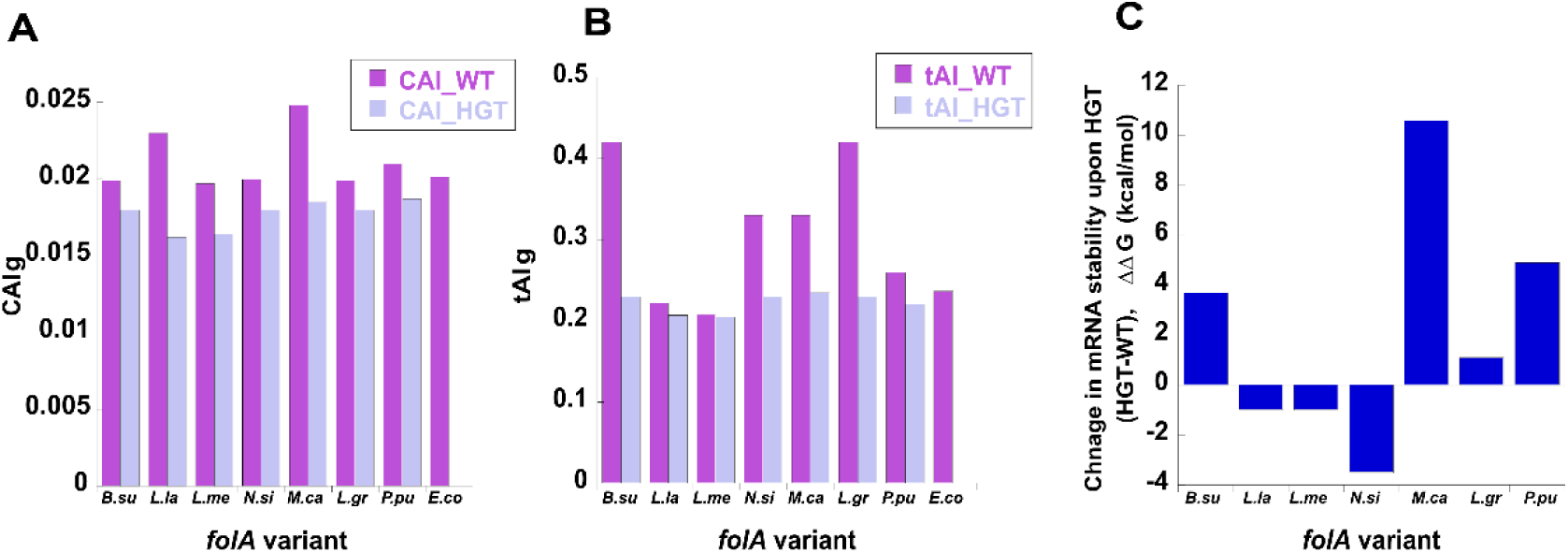
The effect of HGT on codon optimality and 5’-end mRNA stability of the transferred genes. *folA* variants are marked as in **Table 1**. **(A,B)** Change in the (A) CAIg and (B) tAIg values of *folA* genes upon transfer from the original organism (CAI_WT) to *E. coli* (CAI_HGT). **(C)** Change in mRNA folding stability (ΔΔG, kcal/mol) of 30-nt long 5’-end mRNA structure (from nucleotide −25 of the upstream sequence and up to nucleotide +5 within the coding sequence), calculated by subtraction of ΔG in *E. coli* (HGT) from that in the original organism (WT).

**Fig. S2.**
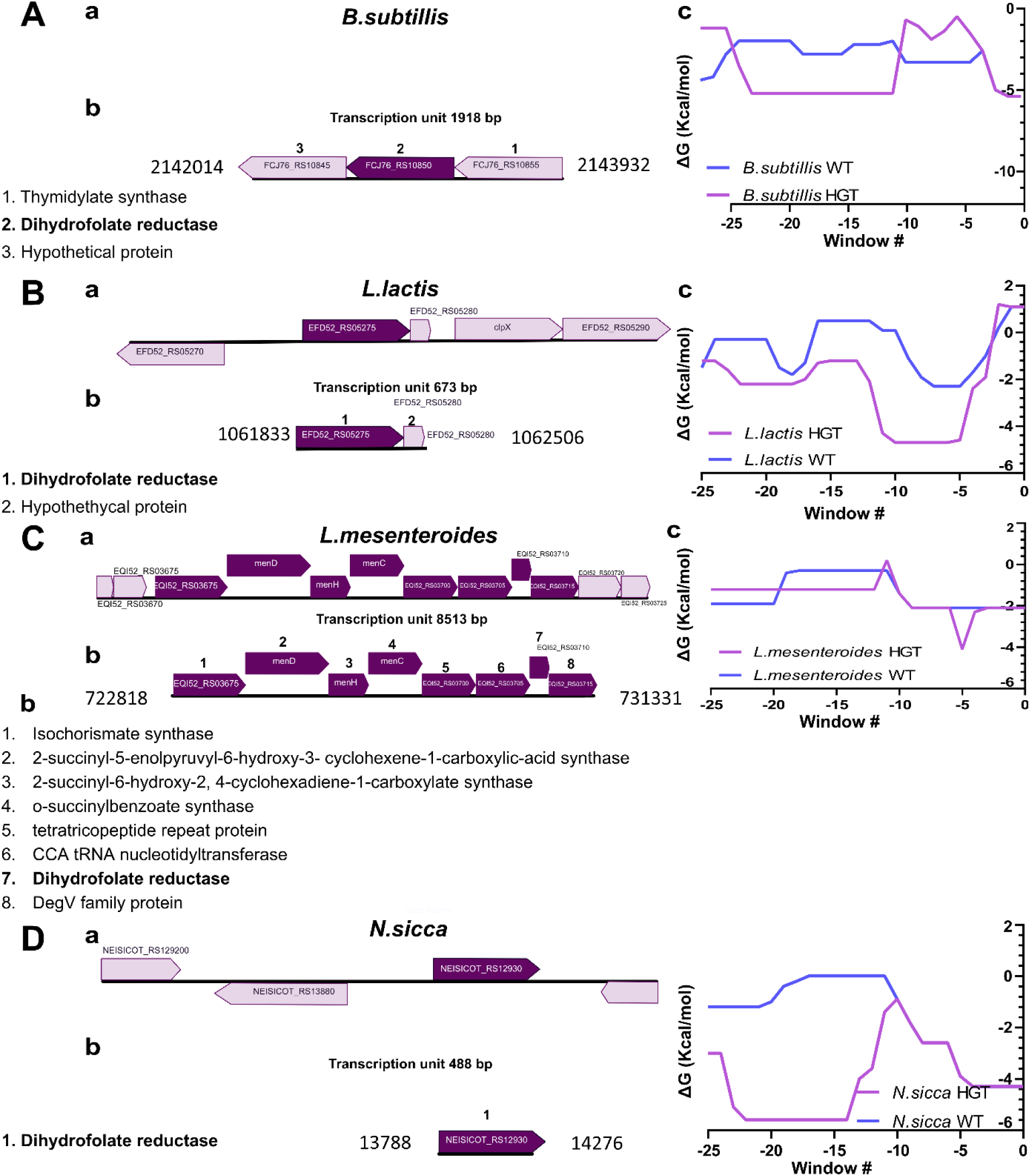
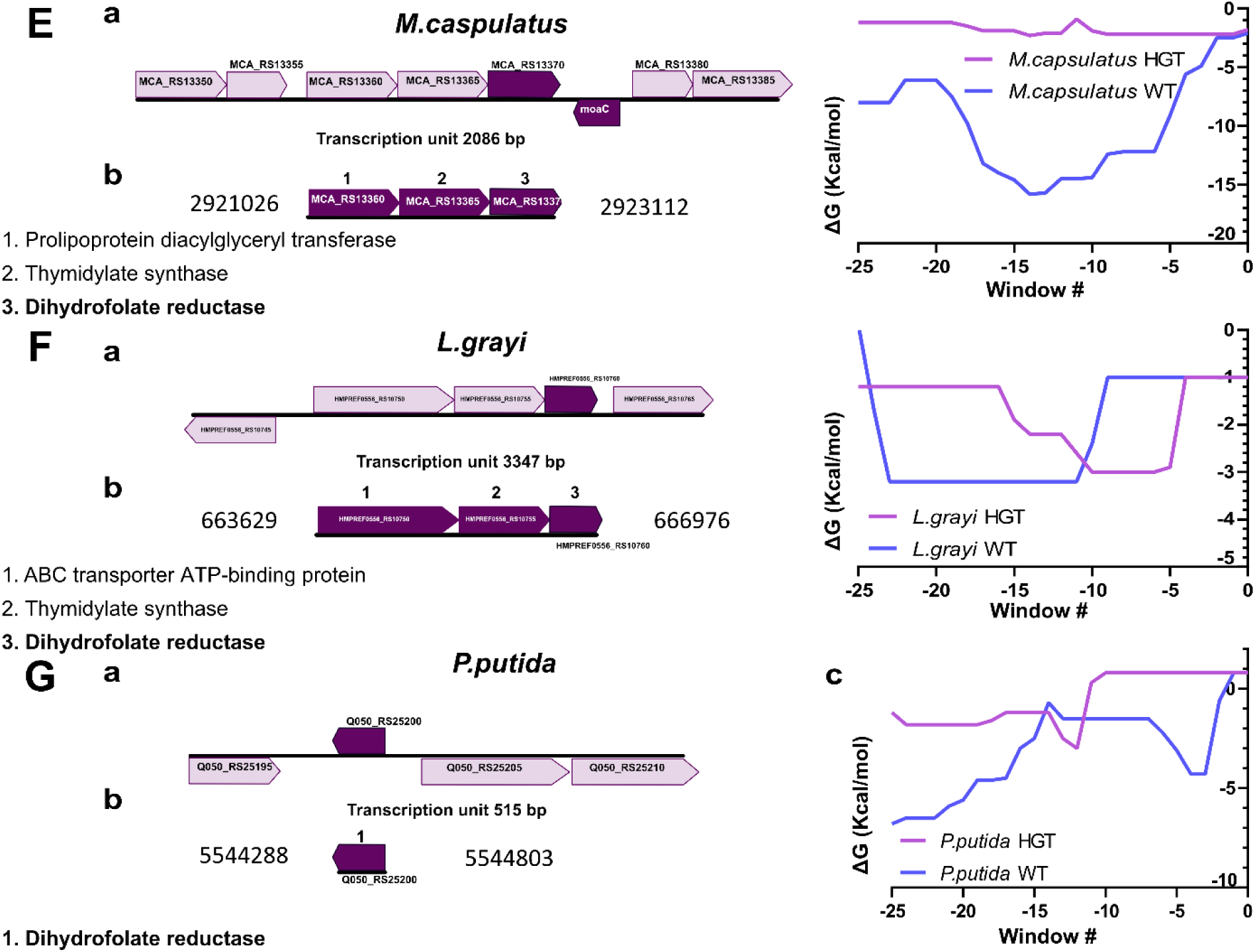
Genomic content/transcript composition of the transferred *folA* genes within the original organisms and change in 5’-end mRNA stability upon HGT to *E. coli.* **(A-G)** transferred *folA* genes. *Left panels.* (a) genomic content and (b) predicted compostion and length of mRNA transcripts. *Right panels*. mRNA folding stability (*D*G, kcal/mol) of transferred *folA* genes in their original transcript content (WT) and upon transfer downstream to *E. coli*’s *folA* endogenous promoter (HGT), calculated using a 30 nucleotide-long sliding window starting at −25 nt upstream to the translation start codon and moving 1 nt at a time for a total of 25 windows.

**Fig. S3.**
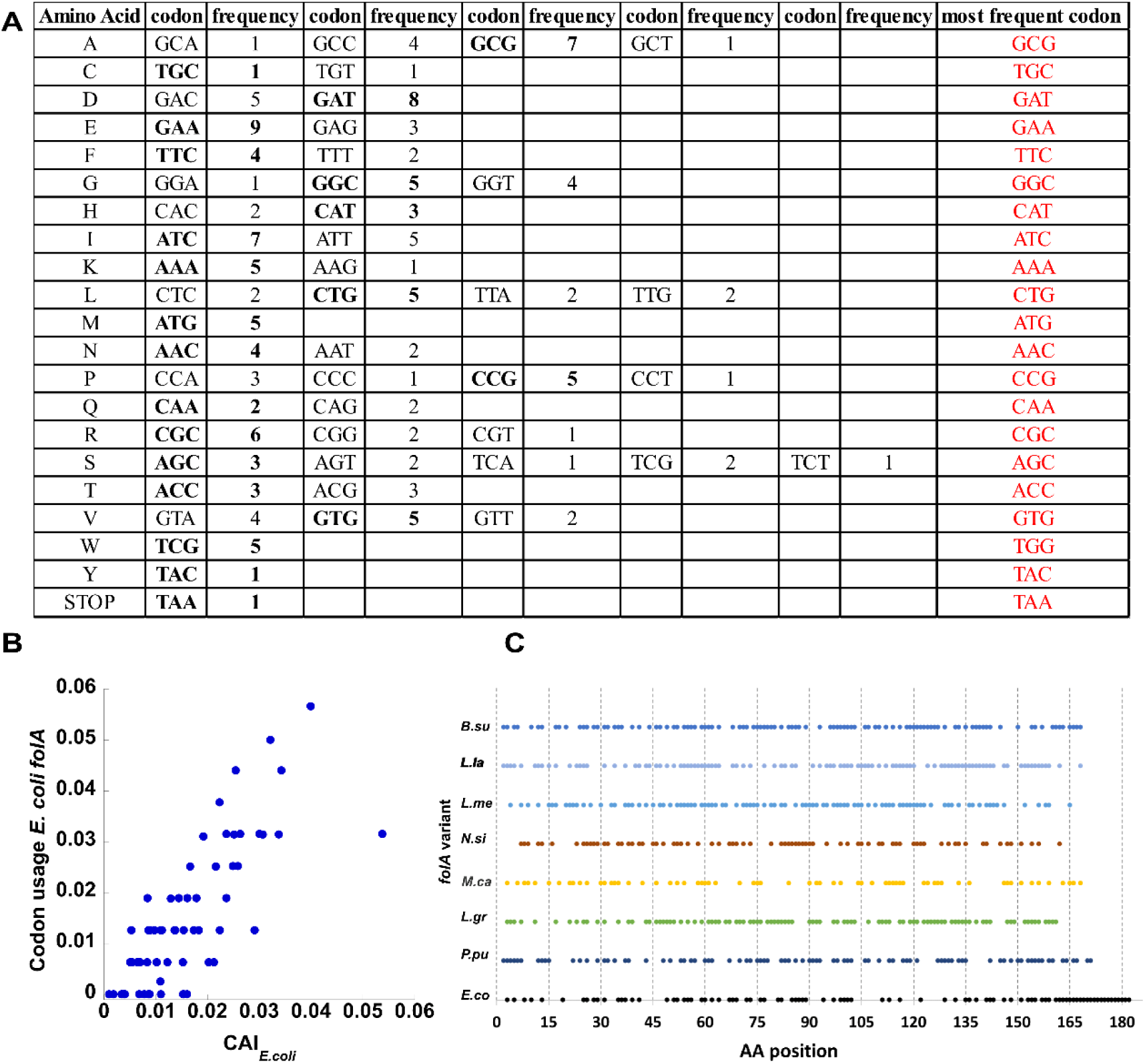
Identification of the most frequent codons in *E. coli*’s *folA* coding sequence. **(A)** Total counts of codons comprising *E. coli*’s *folA* coding sequence and identification of the most freqeunt ones (red). (B) Correlation between the frequencies of each of the codons comprising *E. coli*’s *folA* coding sequence, CAI*_E. coli_* (calculated relatively to the codon composition of the *folA* gene) and the corresponding Codon Adaptation Index (CAI) for each codon as determined for *E. coli*.

**Fig. S4.**
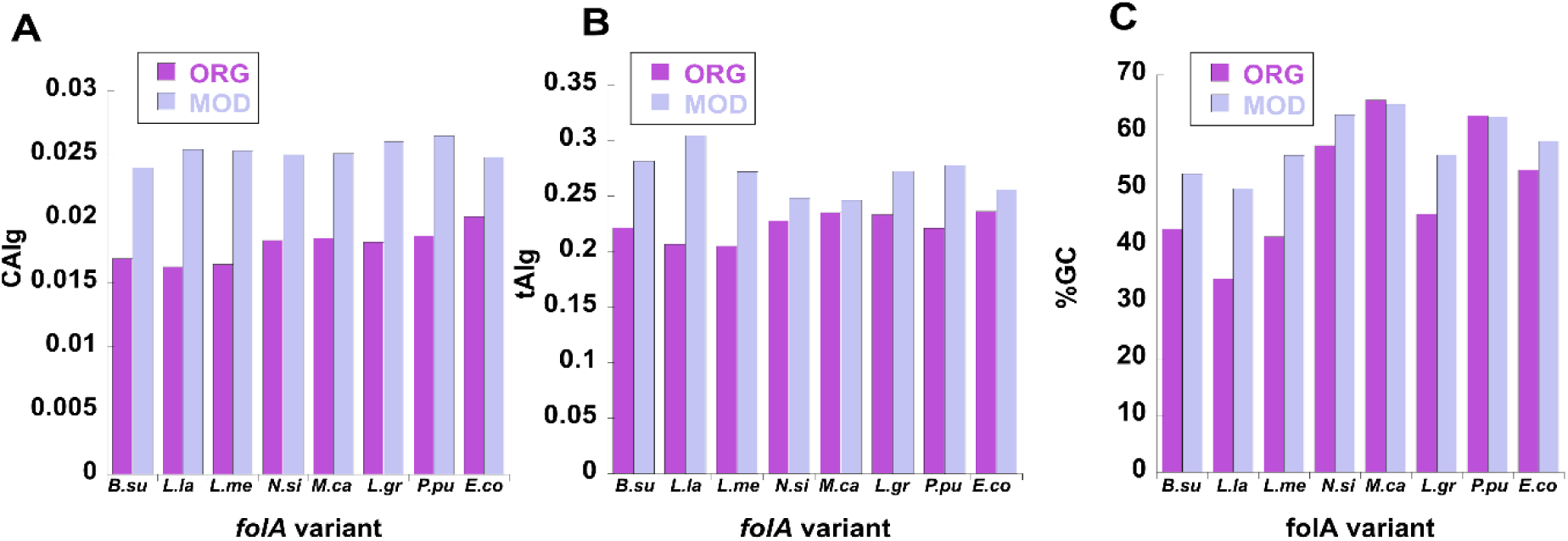
Change in codon optimality and GC content upon original to frequent codon replacement within transferred folA genes. *folA* variants are marked as in **Table 1**. **(A,B)** Change in the (A) CAIg and (B) tAIg values of *folA* genes upon replacemnt of the original (ORG) codons to most frequent (MOD) codons (using *E. coli*’s CAI and tAI metrics, **Table S2**). (C) Change in GC content (%) upon original (ORG) to frequent (MOD) codon replacemnt.

**Fig. S5.**
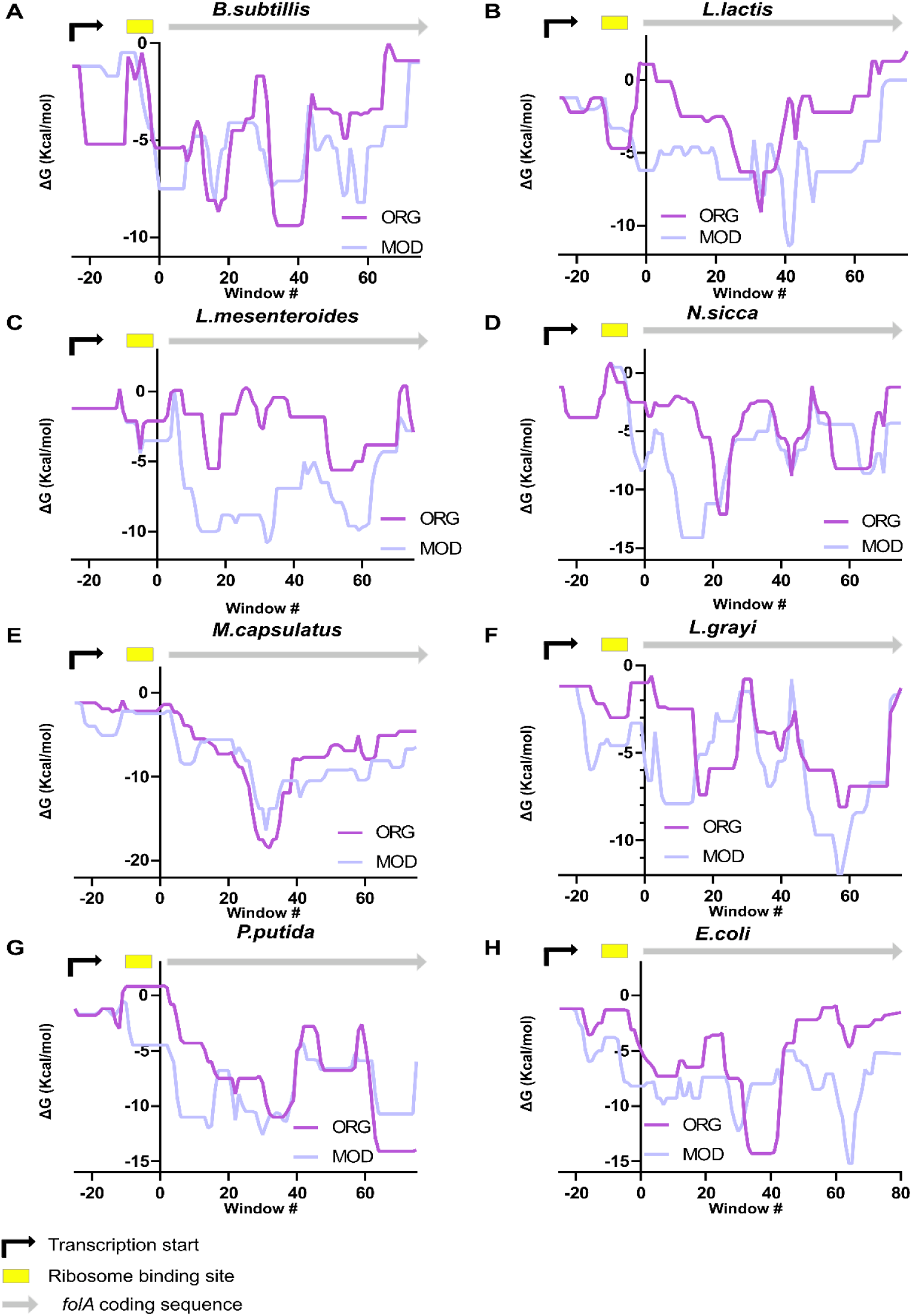
mRNA stability profiles of the xenologous *folA* genes. Stability is expressed in Gibbs free energy change between folded and unfolded states of mRNA transcripts (ΔG, kcal/mol) for the MOD and ORG sequences. Stability is calculated with a 30 nt-long sliding window starting from the transcription start (nucleotide −25 upstream to the translation start codon) in steps of 1 nt.

**Fig. S6.**
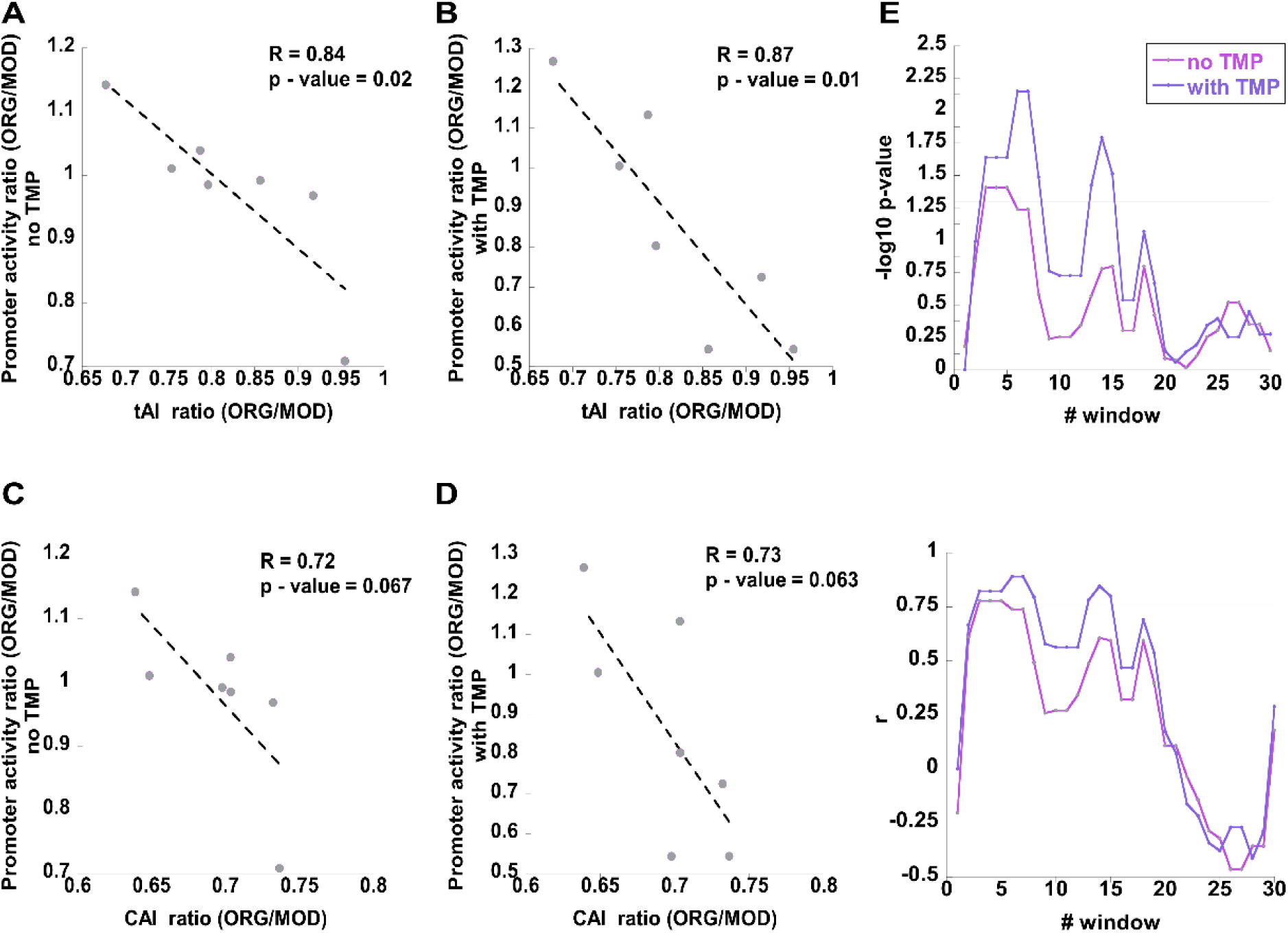
Correlation between changes in promoter activity and changes in codon optimality and mRNA stability upon ORG to MOD codon replacemnt in the absense and presence of IC50 TMP levels. **(A,B)** Changes in promoter activity (calculated as ORG/MOD ratio) are significantly correlated with changes in tAIg values (calculated as ORG/MOD ratio) in the absence (A) and presence (B) of TMP. **(C,D)** Correlation between changes in promoter activity and changes in CAIg values in the absence (A) and presence (B) of TMP are not significant, but follow a trend similar to that observed in (A,B). **(E)** Correlation between the change in promoter activity (calculated as ORG/MOD ratio) and change in mRNA stability (ORG-MOD, ΔΔG, kcal/mol), calculated using a 30 nucleotide-long sliding window starting from the mRNA transcription start (25 nt upstream to the translation start codon) and moving 1 nt at a time for 30 windows, is presented as *p-values* (-log10[p-value]), upper panel, and correlation coefficients, *r*, lower panel, for Spearman non-parametric test. The horizontal bar at the upper panel indicates the significance barrier for *p* < 0.05. *Pink*, no TMP. *Purple*, IC50 levels of TMP.

**Fig. S7.**
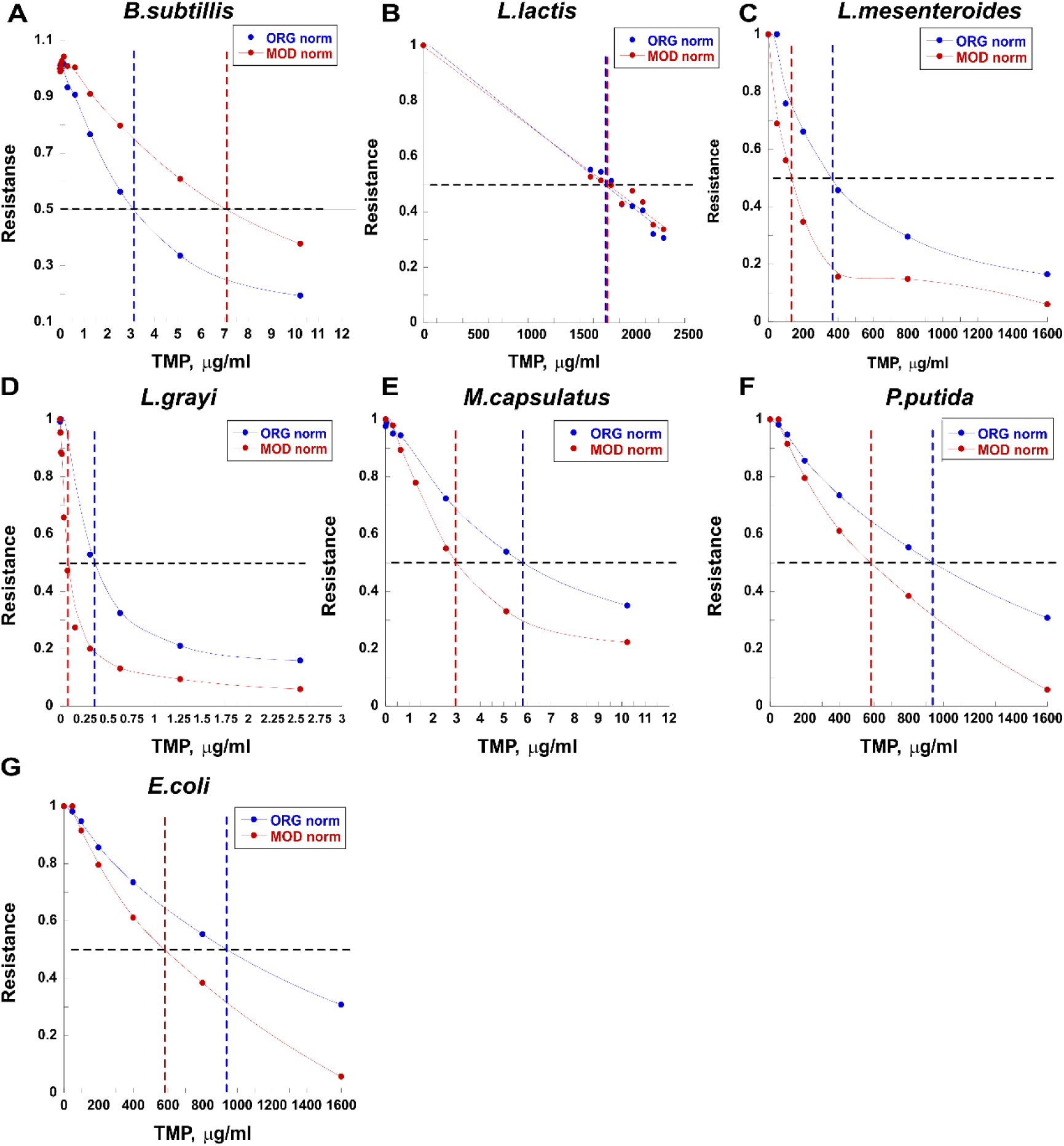
Calculation of TMP IC50 levels for *E. coli* strains carrying ORG and MOD *folA* genes. **(A-G)** Normalized growth of E. coli strains carrying ORG (blue) or MOD (red) *folA* gene is plotted as a function of TMP concentrations. Dashed line intersections with the black line indicate TMP concentrations at which growth of each variant reaches 50% of its growth in the absence of TMP (IC50).

**Fig. S8.**
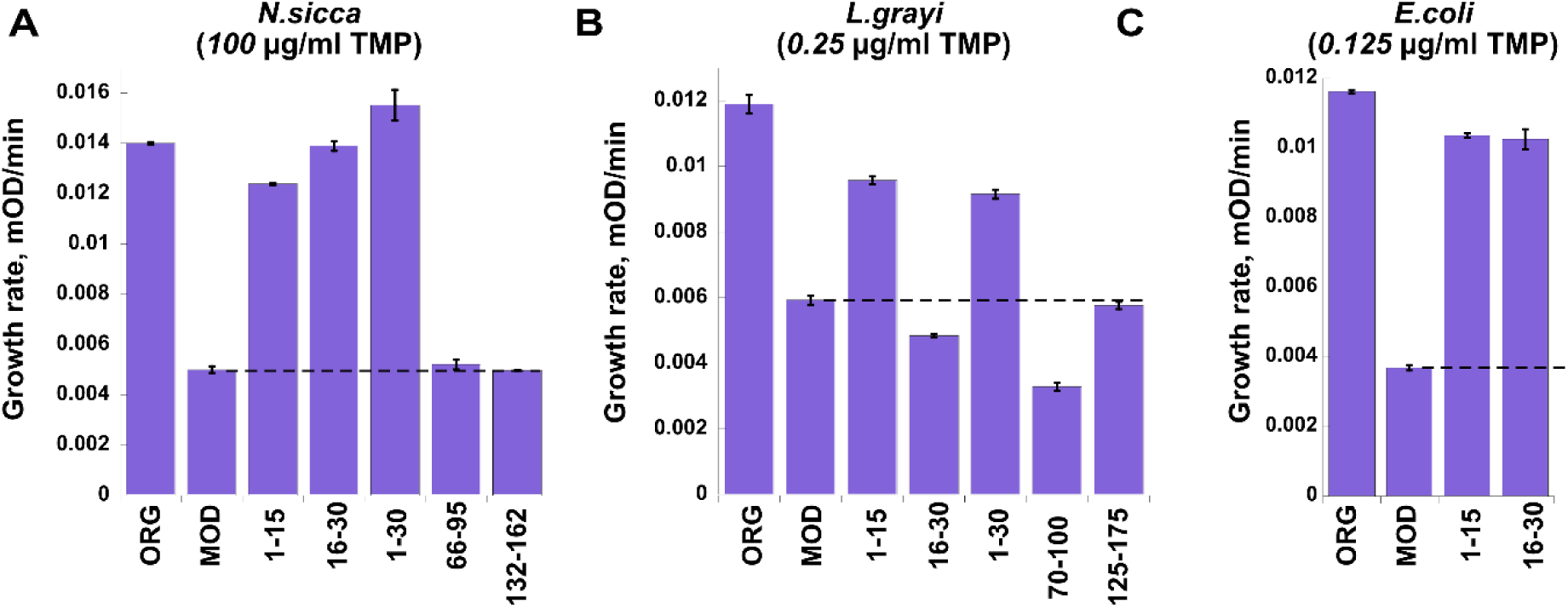
Changes in growth rates of *E. coli* strains carrying modified xenologous and endogenous *folA* genes. Growth rates of strains carrying fully modified (MOD), original (ORG) and chimeric *folA* sequences from **(A)** *N. sicca*, **(B)** *l. grayi*, and **(C)** *E. coli* in the presence of sub-MIC TMP (concentration is shown in parenthesis). Modified codons composing *folA* sequences (MOD) were partially replaced back to original codons by exchanging the modified codons found within codons 1-15, codons 16-30, codons 1-30, middle 30 codons (codons 66-95 in *N. sicca folA*, 70-100 in *L. grayi folA*), last 30 codons (codons 132-162 in *N. sicca folA*, 125-175 in *L. grayi folA*). Dashed lines mark the growth of strains with fully MOD *folA*.

**Fig S9.**
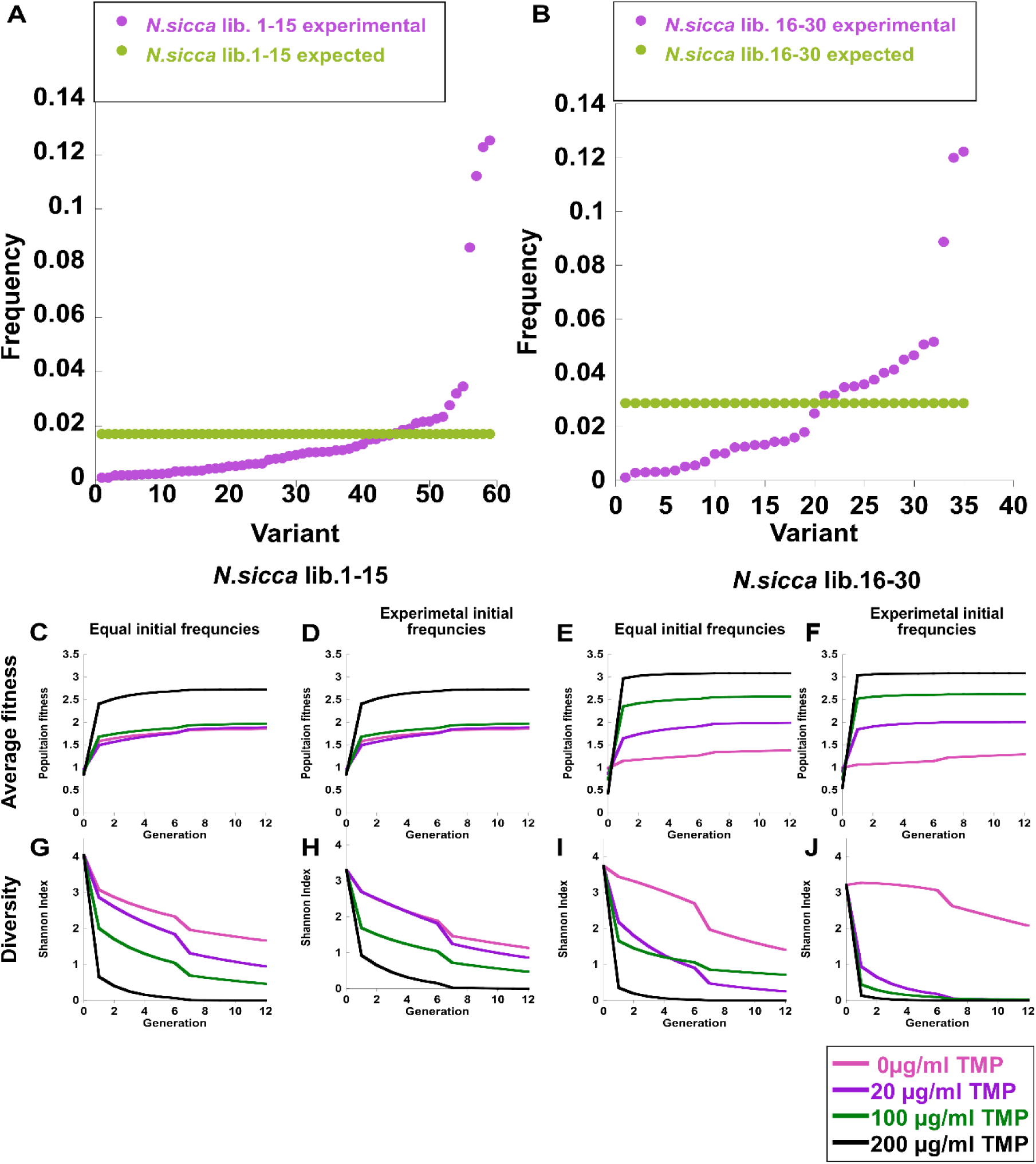
SodaPop simulations of population effects of selection on *N. sicca* libraries 1-15 and 16-30 run with equalized or experimentally defined initial library frequencies. **(A, B)** The expected (equalized, *green*) and experimentally determined (*pink*) initial frequencies of the library variants. **(C-J)** The change in the *N. sicca* lib. 1-15 and 16-30 population average fitness and Shannon diversity simulated with equalized (C, G, E, I) and experimentally determined (D, F, H, G) initial frequencies throughout 12 generation with 0, 20, 100, and 200 μg/ml TMP.

**Fig S10.**
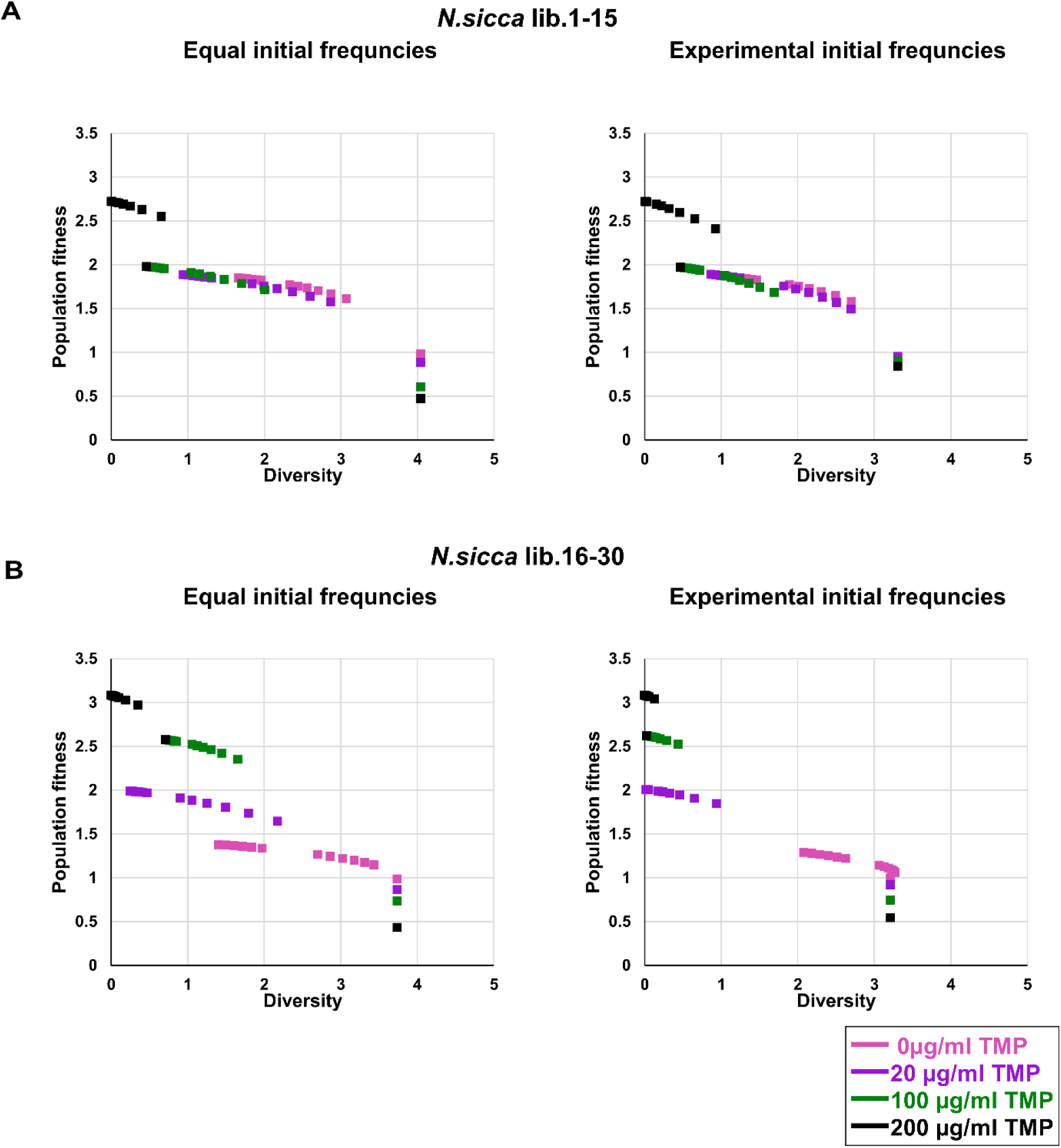
Correlation between the simulated population average fitness and Shannon diversity for N. sicca libraries 1-15 and 16-30. The simulated value for population fitness and diversity was calculated for 12 generations in 4 antibiotic concentrations (0, 20, 100, and 200 µgr/ml TMP). The starting point for each population is the highest diversity and lowest fitness (bottom right corner for each color). As the generations pass, the population average fitness increases and the population Shannon diversity decreases. The rate of this phenomenon corresponds to the slope for each condition (TMP concertation). In general, the higher the antibiotic concentration the faster the increase in the population fitness and drop in diversity. The average fitness of *N. sicca* lib. 1-15 reaches lower values that that of lib. 16-30. This is attributed to the increased clonal interference of the variants. This trend is similar when the initial frequencies are equal or biased, like in the real experiment.

**Fig S11.**
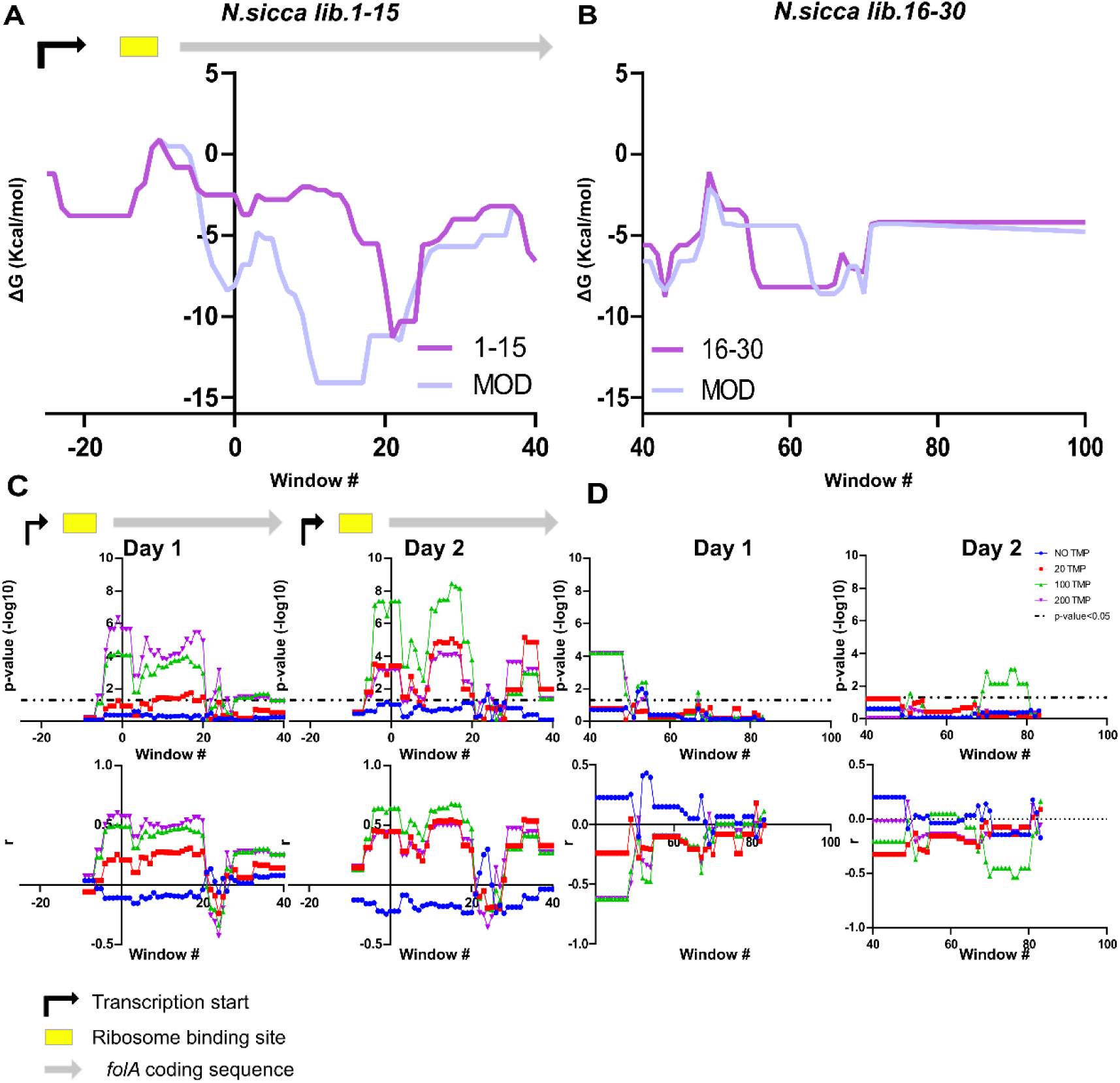
Correlation of fitness effects with mRNA stability in *N. sicca* libraries 1-15 and 16-30 after day1 and day2 of selection at a range of TMP concentrations. mRNA stability was calculated using a 30 nucleotide-long sliding window in steps of 1 nt starting **(A)** from nucleotide −25 and calculated for a modified (MOD) *N. sicca folA* sequence and sequence in which synonymous codons between codons (1-15) were replaced back to the original codons, **(B)** from nucleotide +40 and calculated for a modified (MOD) *N. sicca folA* sequence and sequence in which synonymous codons between codons (16-30) were replaced back to the original codons. **(C, D)** *Upper panels. P-values* for Spearman non-parametric test for association between fitness effects and mRNA stability. *Lower panels.* Corresponding Spearman’s correlation coefficients, *r.*

**Fig S12.**
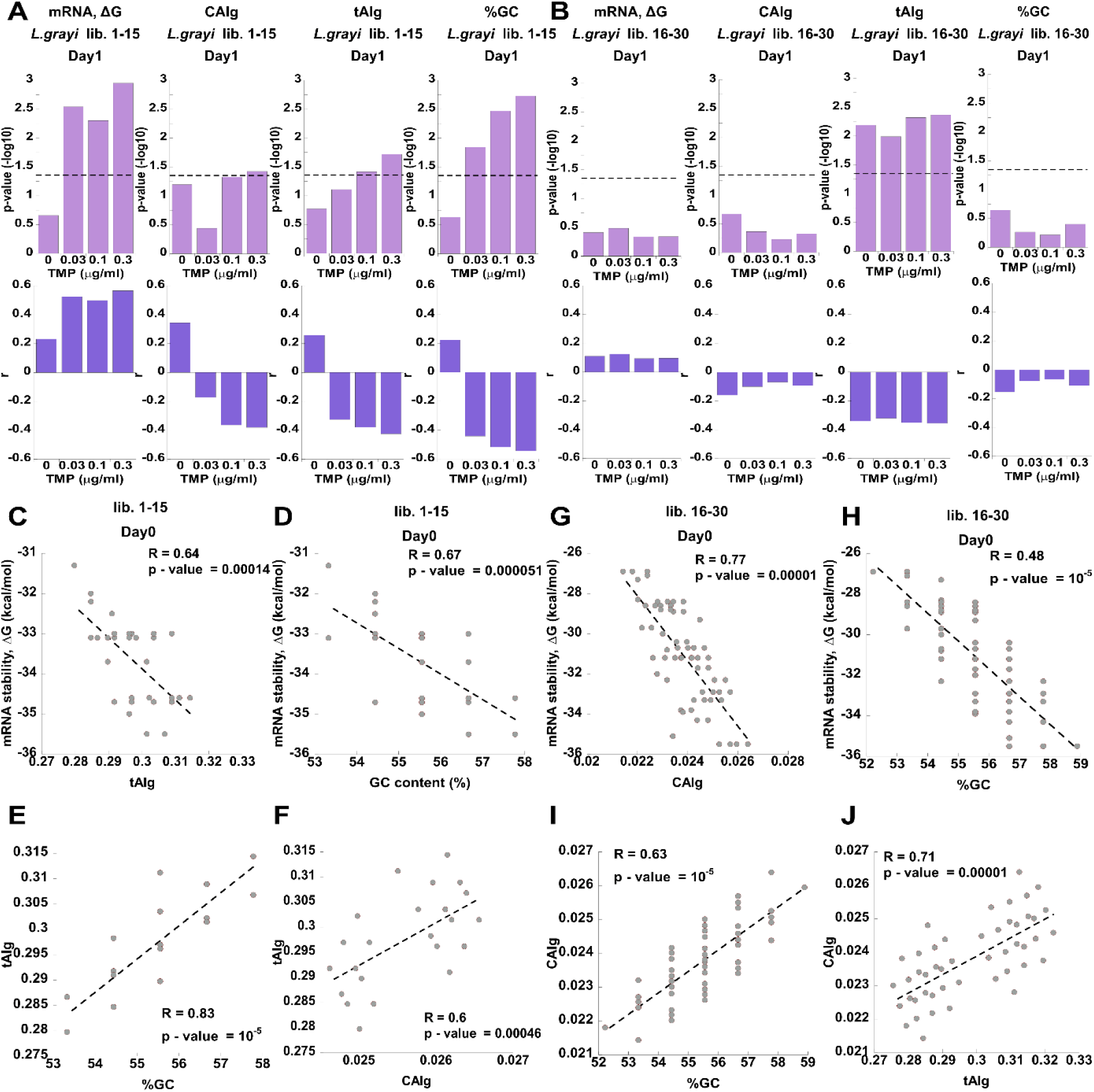
Correlation of fitness effects with *folA* sequence composition in *L. grayi* libraries 1-15 (A) and 16-30 (B) after day1 of selection at a range of TMP concentrations. **(A, B)** *Upper panels. P-values* for Spearman non-parametric test for association between fitness effects and mRNA stability (calculated for a single fragment spanning 114 nucleotides from nucleotide −25, the beginning of transcription, and up to nucleotide +90), codon optimality (CAIg, tAIg values), and GC content. *Lower panels.* Corresponding Spearman’s correlation coefficients, *r.* **C-J**. Pearson’s correlations between mRNA stability, codon optimality (CAIg, tAIg), and GC content for variants comprising naïve libraries (day0). Pearson’s p-values and R values are shown.

**Fig S13.**
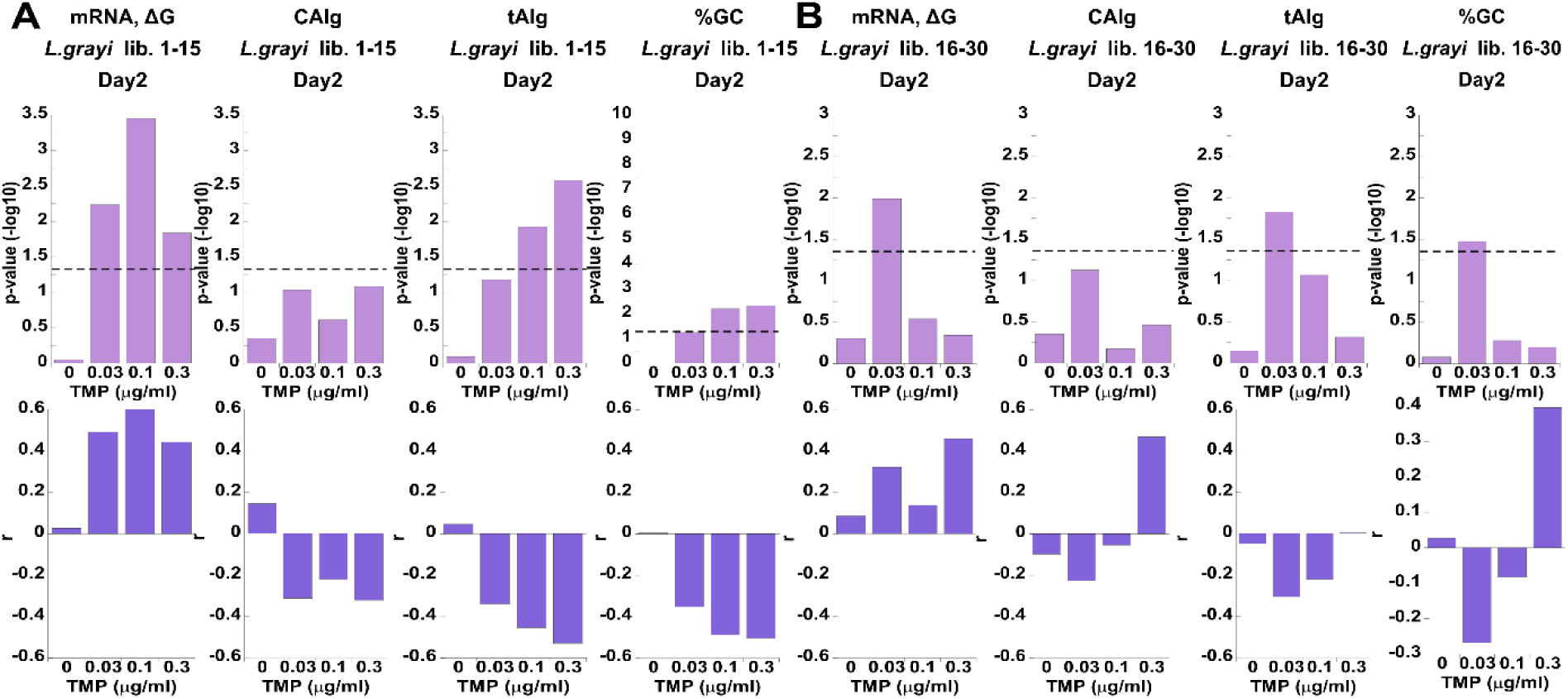
Correlation of fitness effects with *folA* sequence composition in *L. grayi* libraries 1-15 (A) and 16-30 (B) after day2 of selection at a range of TMP concentrations. **(A, B)** *Upper panels. P-values* for Spearman non-parametric test for association between fitness effects and mRNA stability (calculated for a single fragment spanning 114 nucleotides from nucleotide −25, the beginning of transcription, and up to nucleotide +90), codon optimality (CAIg, tAIg values), and GC content. *Lower panels.* Corresponding Spearman’s correlation coefficients, *r.*

**Fig S14.**
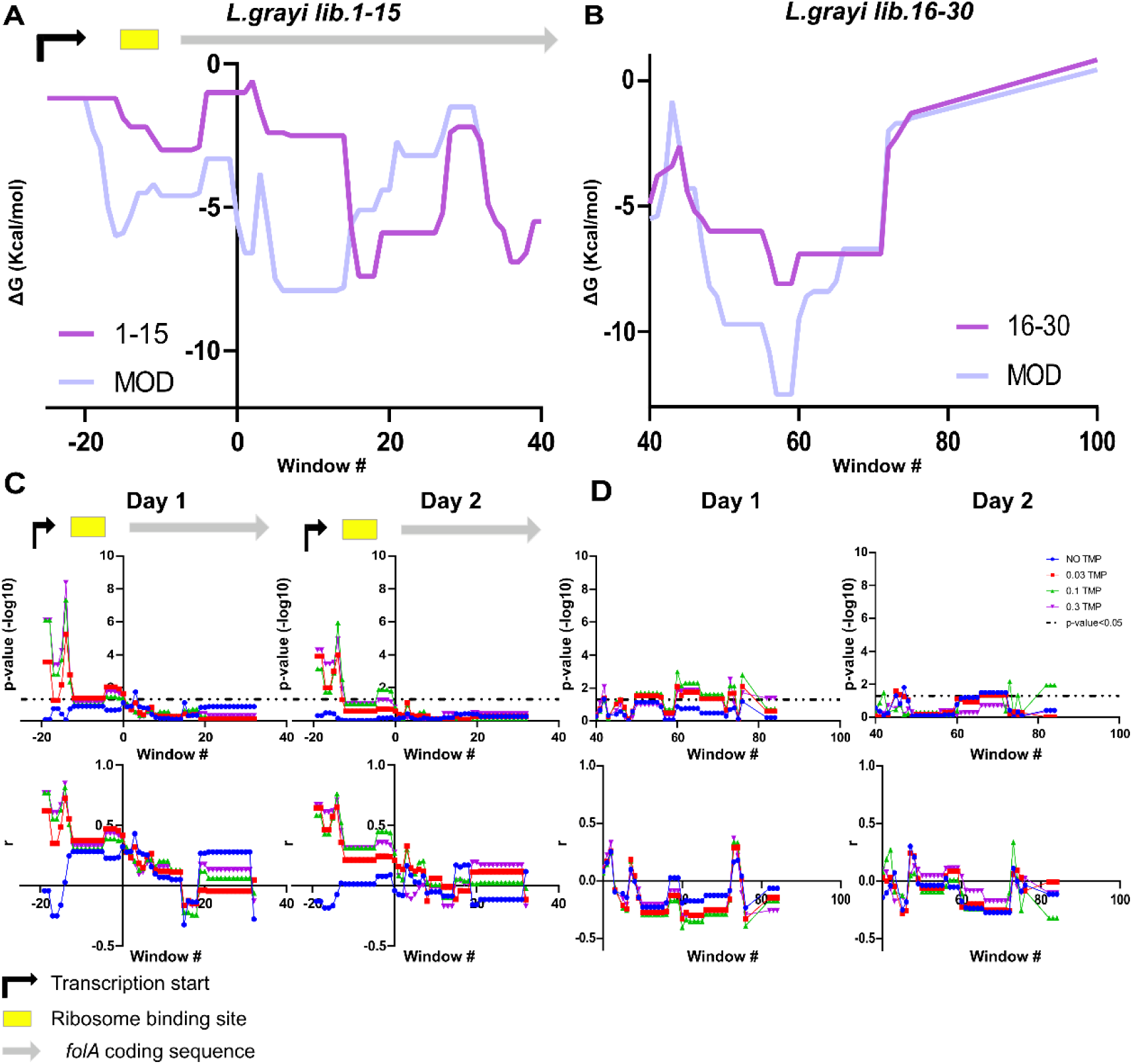
Correlation of fitness effects with mRNA stability in *L. grayi* libraries 1-15 and 16-30 after day1 and day2 of selection at a range of TMP concentrations. mRNA stability was calculated using a 30 nucleotide-long sliding window in steps of 1 nt starting **(A)** from nucleotide −25 and calculated for a modified (MOD) *L. grayi folA* sequence and sequence in which synonymous codons between codons (1-15) were replaced back to the original codons, **(B)** from nucleotide +40 and calculated for a modified (MOD) *N. sicca folA* sequence and sequence in which synonymous codons between codons (16-30) were replaced back to the original codons. **(C, D)** *Upper panels. P-values* for Spearman non-parametric test for association between fitness effects and mRNA stability. *Lower panels.* Corresponding Spearman’s correlation coefficients, *r.*

**Fig S15.**
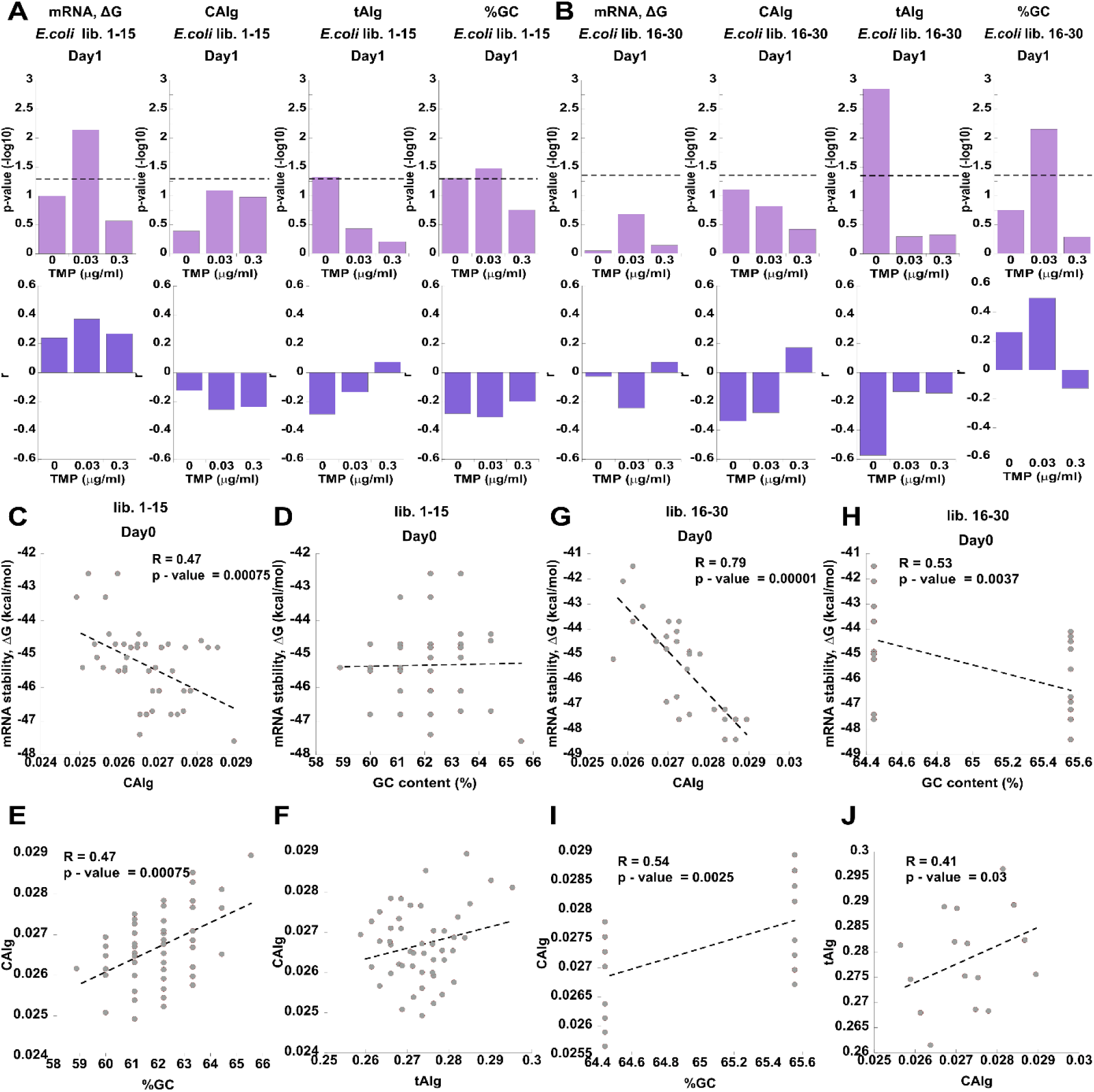
Correlation of fitness effects with *folA* sequence composition in *E. coli* libraries 1-15 (A) and 16-30 (B) after day1 of selection at a range of TMP concentrations. **(A, B)** *Upper panels. P-values* for Spearman non-parametric test for association between fitness effects and mRNA stability (calculated for a single fragment spanning 114 nucleotides from nucleotide −25, the beginning of transcription, and up to nucleotide +90), codon optimality (CAIg, tAIg values), and GC content. *Lower panels.* Corresponding Spearman’s correlation coefficients, *r.* **C-J**. Pearson’s correlations between mRNA stability, codon optimality (CAIg, tAIg), and GC content for variants comprising naïve libraries (day0). Pearson’s p-values and R values are shown.

**Fig S16.**
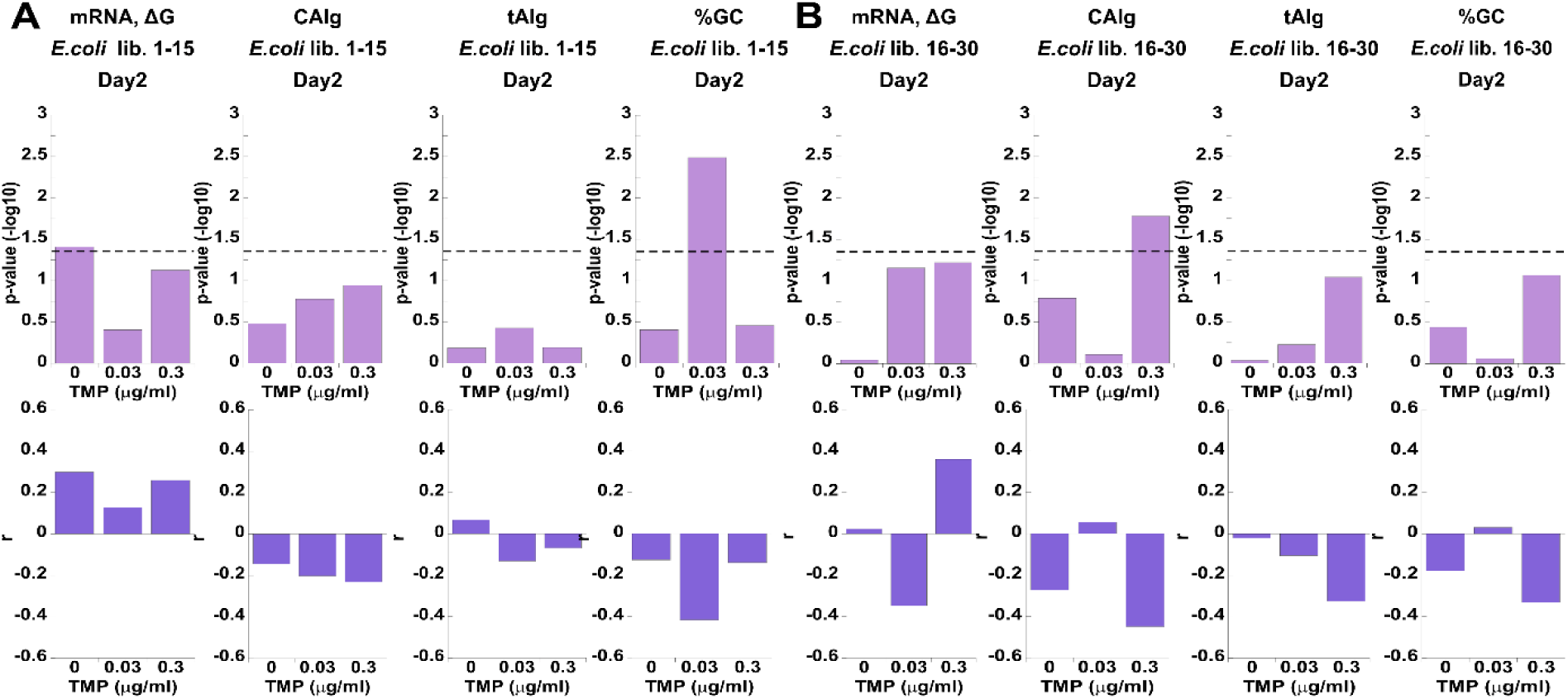
Correlation of fitness effects with *folA* sequence composition in *E. coli* libraries 1-15 (A) and 16-30 (B) after day2 of selection at a range of TMP concentrations. **(A, B)** *Upper panels. P-values* for Spearman non-parametric test for association between fitness effects and mRNA stability (calculated for a single fragment spanning 114 nucleotides from nucleotide −25, the beginning of transcription, and up to nucleotide +90), codon optimality (CAIg, tAIg values), and GC content. *Lower panels.* Corresponding Spearman’s correlation coefficients, *r.*

**Fig S17.**
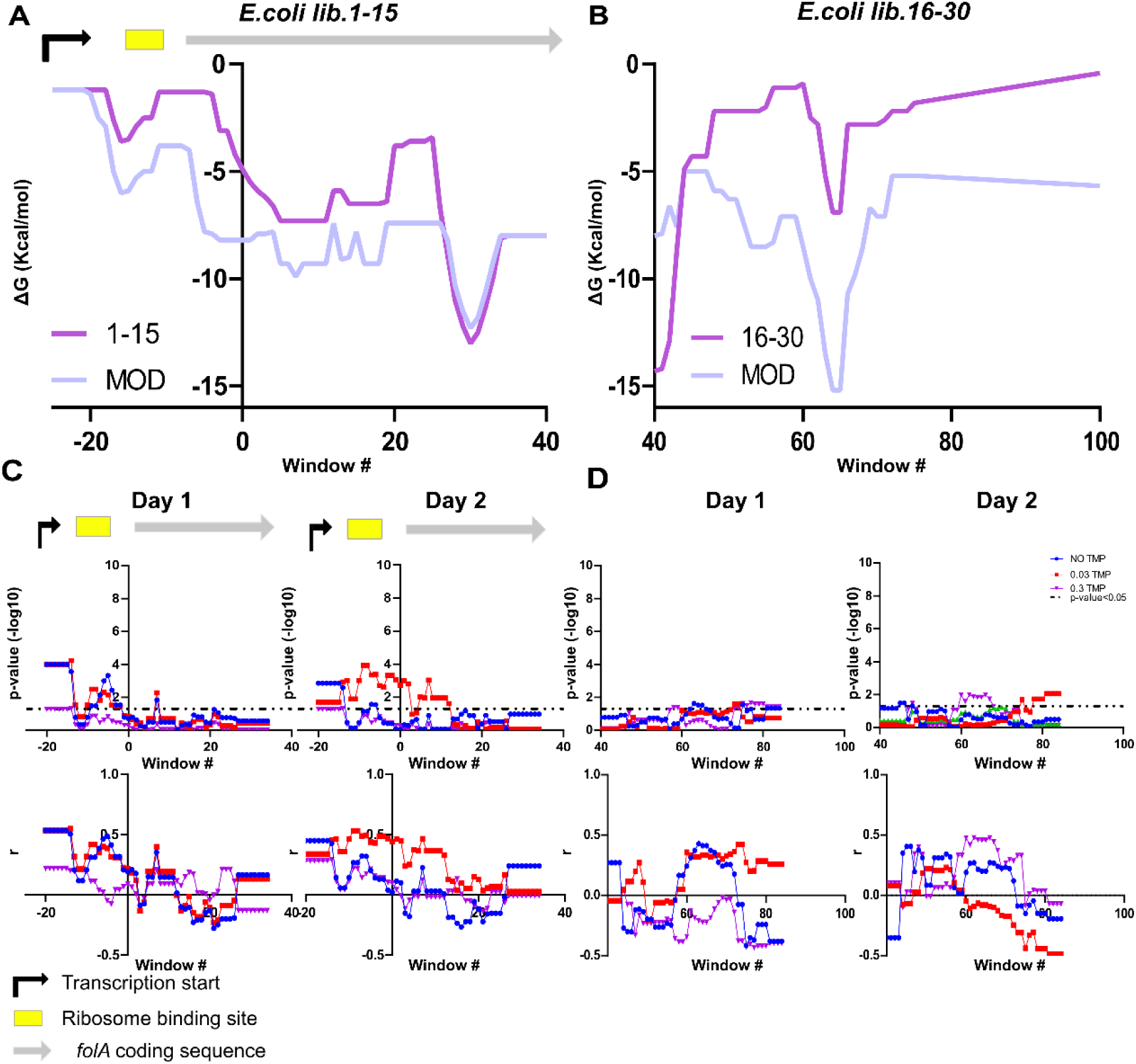
Correlation of fitness effects with mRNA stability in *E. coli.* libraries 1-15 and 16-30 after day1 and day2 of selection at a range of TMP concentrations. mRNA stability was calculated using a 30 nucleotide-long sliding window in steps of 1 nt starting **(A)** from nucleotide −25 and calculated for a modified (MOD) *E. coli folA* sequence and sequence in which synonymous codons between codons (1-15) were replaced back to the original codons, and **(B)** from from nucleotide +40 and calculated for a modified (MOD) *E. coli folA* sequence and sequence in which synonymous codons between codons (16-30) were replaced back to the original codons. **(C, D)** *Upper panels. P-values* for Spearman non-parametric test for association between fitness effects and mRNA stability. *Lower panels.* Corresponding Spearman’s correlation coefficients, *r.*

**Fig S18.**
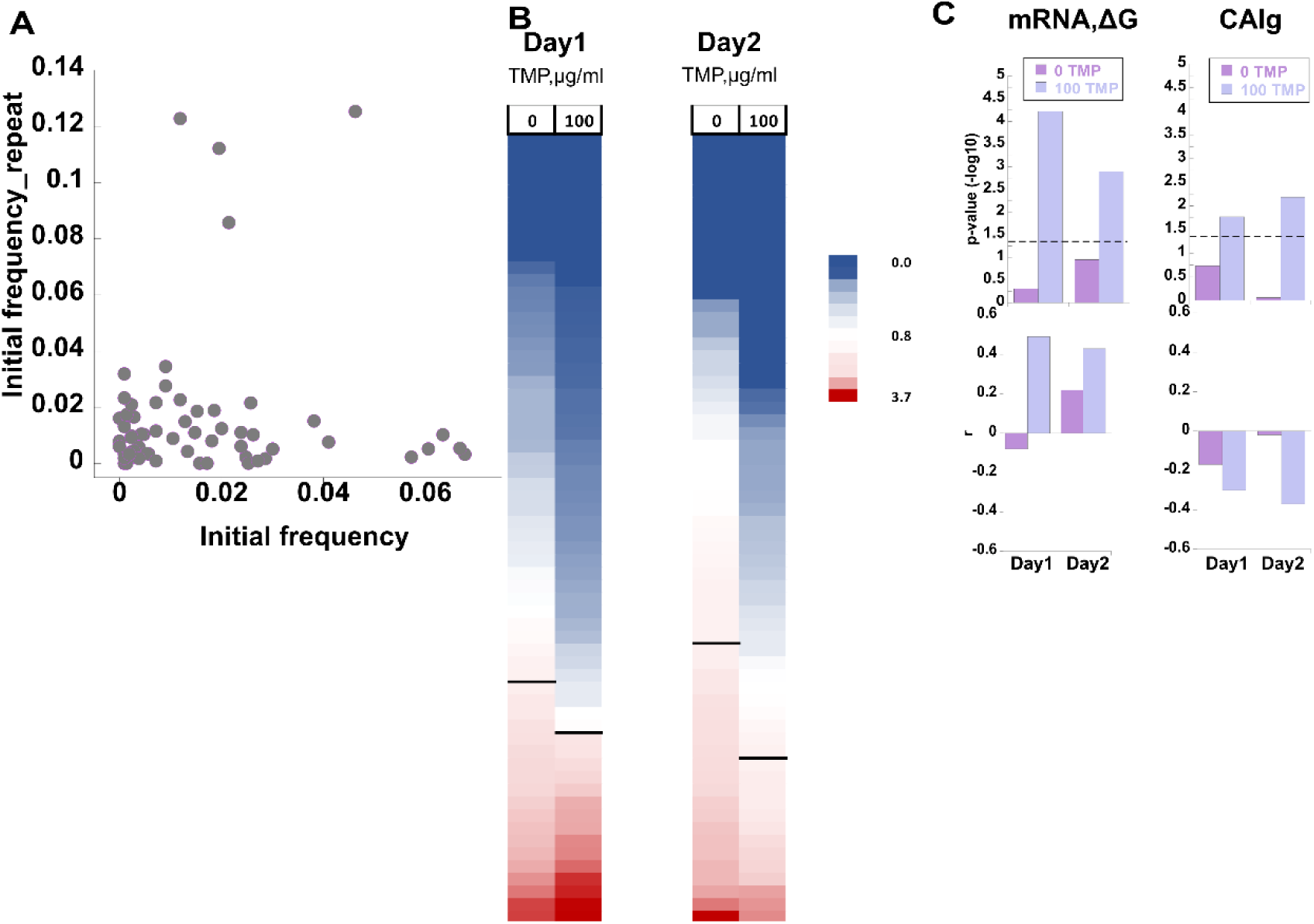
Repeat of selection on *N. sicca* lib. 1-15 after the perturbation in the initial variant frequencies. **(A)** Correlation between the initial frequencies prior to (Initial frequency) and after (Initial frequency_repeat) introduction of the perturbation. **(B)** Relative fitness of the individual library variants was sorted from lowest (*blue*) to highest (*red*) independently at each selection regime (no TMP and 100 μg/ml TMP) and presented as heat maps. (Note that variants at each row are not necessarily identical, as their fitness may change upon change in the selection regime). Black bars separate variants whose fitness dropped upon selection (fitness < 1) from those whose fitness has increased (fitness > 1). The heat map ruler indicates the fold change in the relative fitness of the individual variants. **(C)** *Upper panels. P-values* for Spearman non-parametric test for the association between the fitness effects and (*A*) mRNA stability (calculated for a single fragment spanning 114 nucleotides from nucleotide −25, the beginning of transcription, and up to nucleotide +90), and fitness effects and (*B*) codon optimality (CAIg). *Lower panels.* Corresponding Spearman’s correlation coefficients, *r.*

**Fig S19.**
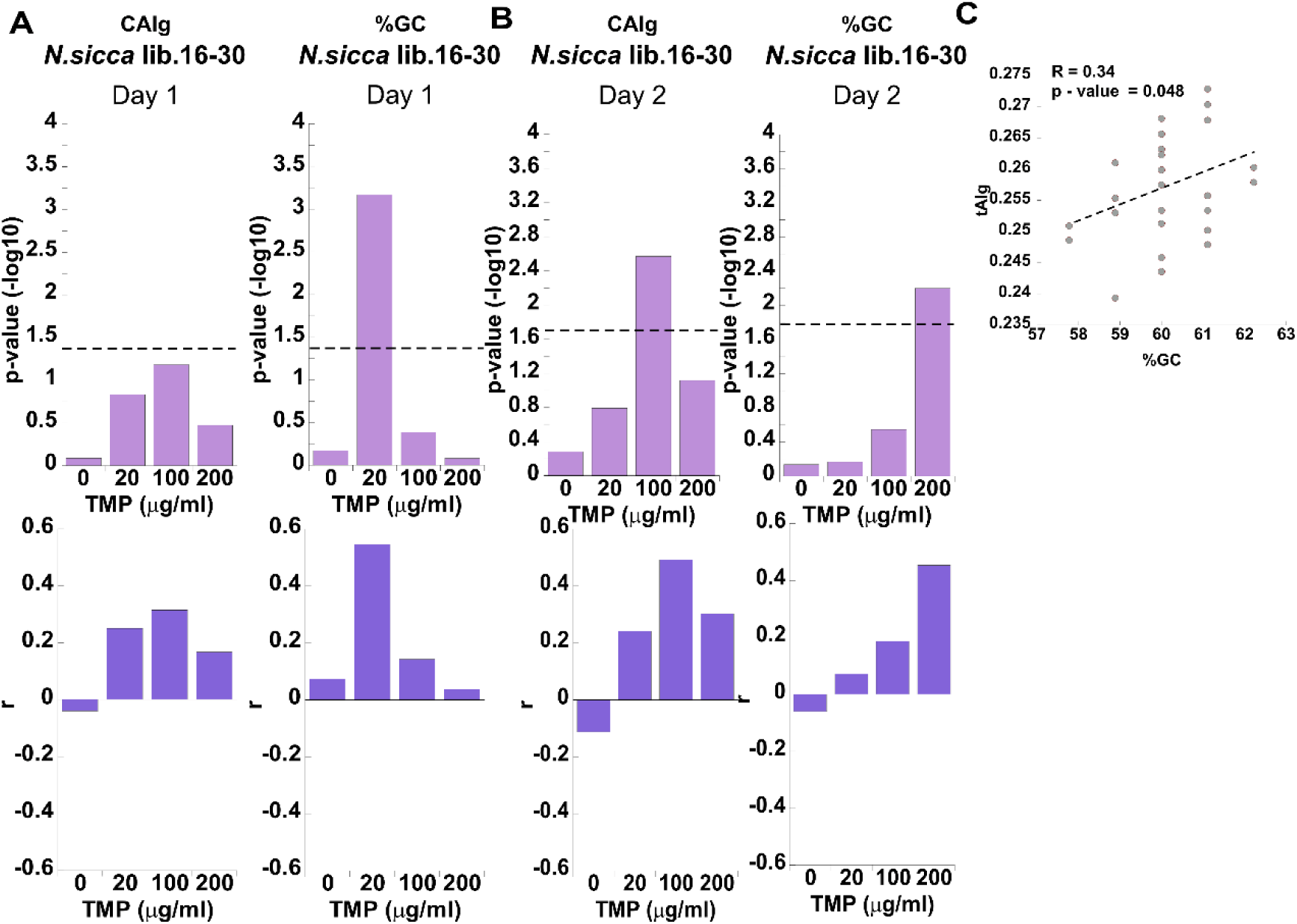
Correlation of fitness effects with *folA* sequence composition in *N. sicca* library 16-30 on day1 and day2 of selection at a range of TMP concentrations. **(A, B)** *Upper panels. P-values* for Spearman non-parametric test for association between fitness effects and codon optimality (CAIg values), and GC content. *Lower panels.* Corresponding Spearman’s correlation coefficients, *r.* (C) Pearson’s correlations between codon optimality (tAIg) and GC content for variants comprising the naïve libraries (day0). Pearson’s p-value and R are shown.

**Fig S20.**
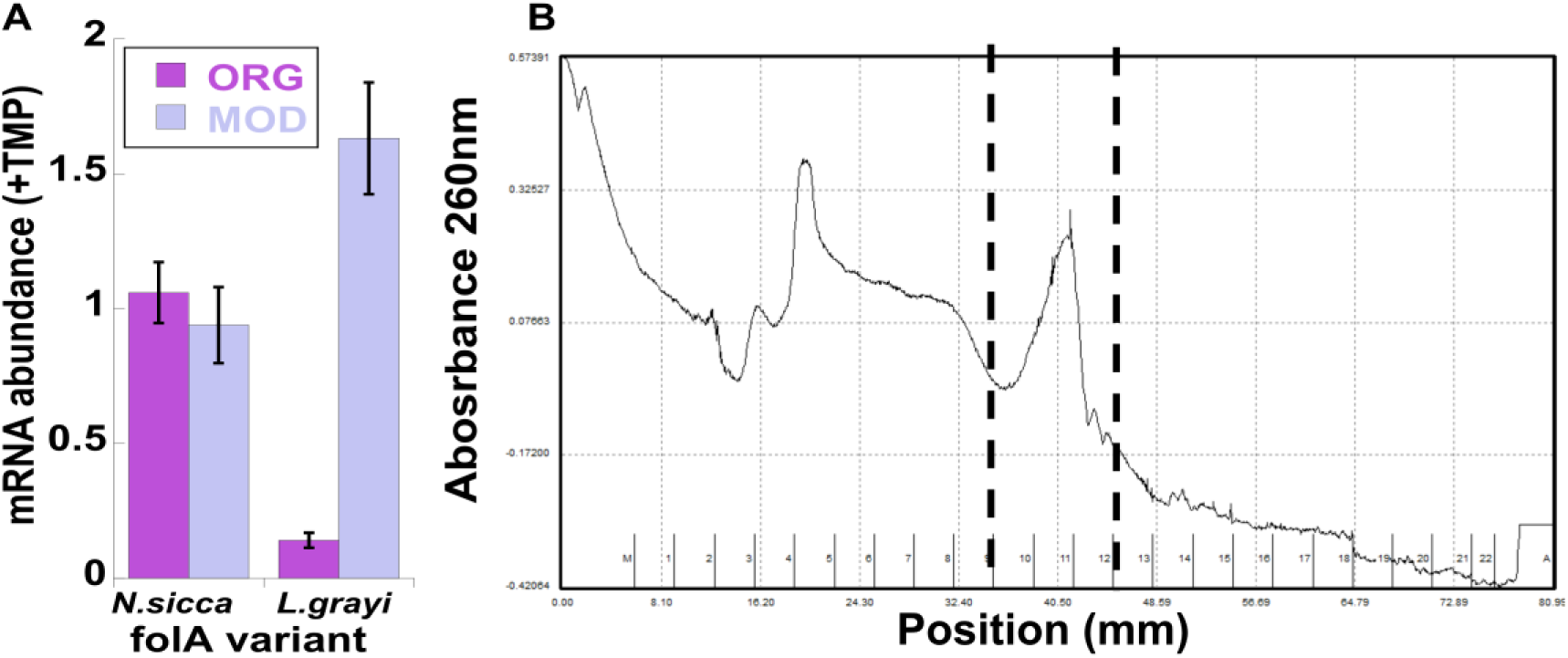
**(A)** The effect of codon usage on mRNA abundance in the presence of IC50 TMP concentrations. (B) A representative polysomal profile. The collected fraction is indicated within the dashed lines.

